# Single cell sequencing of pig lungs reveals immune responses underlying influenza infection and oseltamivir therapy

**DOI:** 10.1101/2024.03.05.581049

**Authors:** Darling Melany de Carvalho Madrid, Weihong Gu, Wesley C. Warren, John P. Driver

**Author notes:** Address correspondence to Dr. John P. Driver, Bond Life Sciences Center Building, 1201 Rollins St, Columbia MO 65201.

## Abstract

Despite pigs being an important species in influenza A virus (IAV) epidemiology and a reliable model of human IAV infections, many aspects of the porcine pulmonary immune system remain poorly understood. Here, we characterized the single cell landscape of lung leukocytes of healthy pigs and then compared them to pigs infected with 2009 pandemic H1N1 IAV with or without oseltamivir antiviral therapy. Our data show conserved features as well as species-specific differences in cell types and cell states compared to human and mouse lung leukocytes. IAV infection induced a robust antiviral transcriptional response in multiple lymphoid and myeloid cell types, as well as distinct patterns of cell-cell cross talk. Oseltamivir treatment reduced these responses. Together our findings describe key events in the pulmonary anti-IAV response of pigs that open new avenues to develop IAV vaccines and therapies. They should also enable the better use of pigs as a model for human IAV infection and immunity.

## INTRODUCTION

The lung is a complex organ composed of the pulmonary endothelium, a layer of squamous endothelial cells lining the entire pulmonary circulation, and a permeable layer of epithelial cells that allows for capillary gas exchange (1). Immune cells dispersed throughout the airway lumen, just beneath the epithelial barrier, surveil the respiratory tract for microorganisms and respond to environmental cues released by the lung’s structural cells. The respiratory immune repertoire is composed of over 20 types of specialized cell types (2), some of which can organize into structures that resemble lymph-nodes following infection (3,4). It also includes cell types that restore epithelial integrity after infection or lung damage (5,6). The primary function of the pulmonary immune system is to prevent microbial pathogens from invading the airway tissues. However, it is essential that such immune responses do not damage the delicate anatomical structure of the lung tissue. Hence, the pulmonary immune system must strike a balance between defending the airway from dangerous infections and avoiding overreaction to commensal bacteria and environmental antigens present at the epithelial boundary.

Pigs are considered an excellent model for studying the human respiratory system since porcine and human lung anatomy and surface area are similar (7). Furthermore, practices such as bronchoscopy and endotracheal imaging can be performed on pigs using human instruments (7,8), which makes swine useful for testing lung-related procedures like anesthetics, respirator intubation, and lung transplantation techniques (9,10). Pigs are also subject to genetic editing, including to prevent severe rejection in human xenotransplantation. As a result, pigs are a possible future source of organs, including lungs (11). Similarities between the pulmonary immune system of pigs and humans support the use of pigs as a biomedical model of human immune-related respiratory diseases (12,13). In this regard, pigs have been used for decades to model pulmonary hypersensitivity reactions in humans (14). Another major focus has been the study of respiratory pathogens in swine. Since pigs and humans are susceptible to many of the same infectious agents, swine are a valuable preclinical model to develop therapeutics and vaccines against human respiratory pathogens. A prime example is influenza A virus (IAV) infections which causes similar pathology and clinical manifestations in pigs and humans (15,16).

Influenza is a significant public health concern that results in mortality and morbidity among humans on a global scale (17,18). In addition, acute respiratory illness caused by influenza virus infections in pigs results in huge financial losses and significant losses for the swine industry (19). Pigs are important in the field of IAV epidemiology due to their ability to facilitate the replication of IAVs that originate in swine, birds, and humans (20). Pigs can therefore serve as a reservoir for IAVs that infect numerous species including humans. Moreover, swine occasionally generate novel IAVs that have the potential to cause pandemics in humans (21). In order to mitigate the health consequences of IAV for humans through improved utilization of pigs as models for human IAV infections and by restricting IAV access to swine herds, it is imperative to comprehend the host defense mechanisms that regulate viral replication and pathogenesis in pigs.

Here we used single-cell RNA sequencing (scRNA-seq) to create a cell atlas of lung leukocytes from newly weaned infant pigs. We compared our data to human and mouse lung scRNA-seq datasets to identify similarities and differences in immune cell populations and transcriptional profiles (2,22). Additionally, we compared lung leukocytes between (i) healthy pigs, (ii) pigs infected with 2009 pandemic H1N1 influenza virus (pdmH1N1), and (iii) pdmH1N1 infected pigs treated with the neuraminidase inhibitor oseltamivir. Individual cellular transcriptomes were analyzed to determine how IAV infection and antiviral therapy affected lung leukocytes. This single cell profile of the pig lung provides a resource to gain deeper insight into the workings of their pulmonary immune system. These data also comparatively describe how mechanistic modeling from pig, mouse, and human lung immune cells data, can establish the suitability of pigs as a model for studying human pulmonary immune responses.

## RESULTS

### Cellular composition of porcine lung leukocytes

Single-cell RNA-sequencing was performed on the lungs of two healthy (Healthy) five-week-old mixed breed pigs. After removing cells with unusual gene counts and high mitochondrial gene expression, we performed an unsupervised clustering analysis using Seurat (v4.3.0). A total of 19,994 cells led to an integrated set of 15 transcriptionally distinct clusters (clusters 1-15) (Figure 1A), which after using established immune lineage markers (Figure 1B) annotated populations within myeloid lineages, such as dendritic cells (DC), monocytes, and macrophages (see markers used in Figure 1C) were apparent.

**Figure 1.**
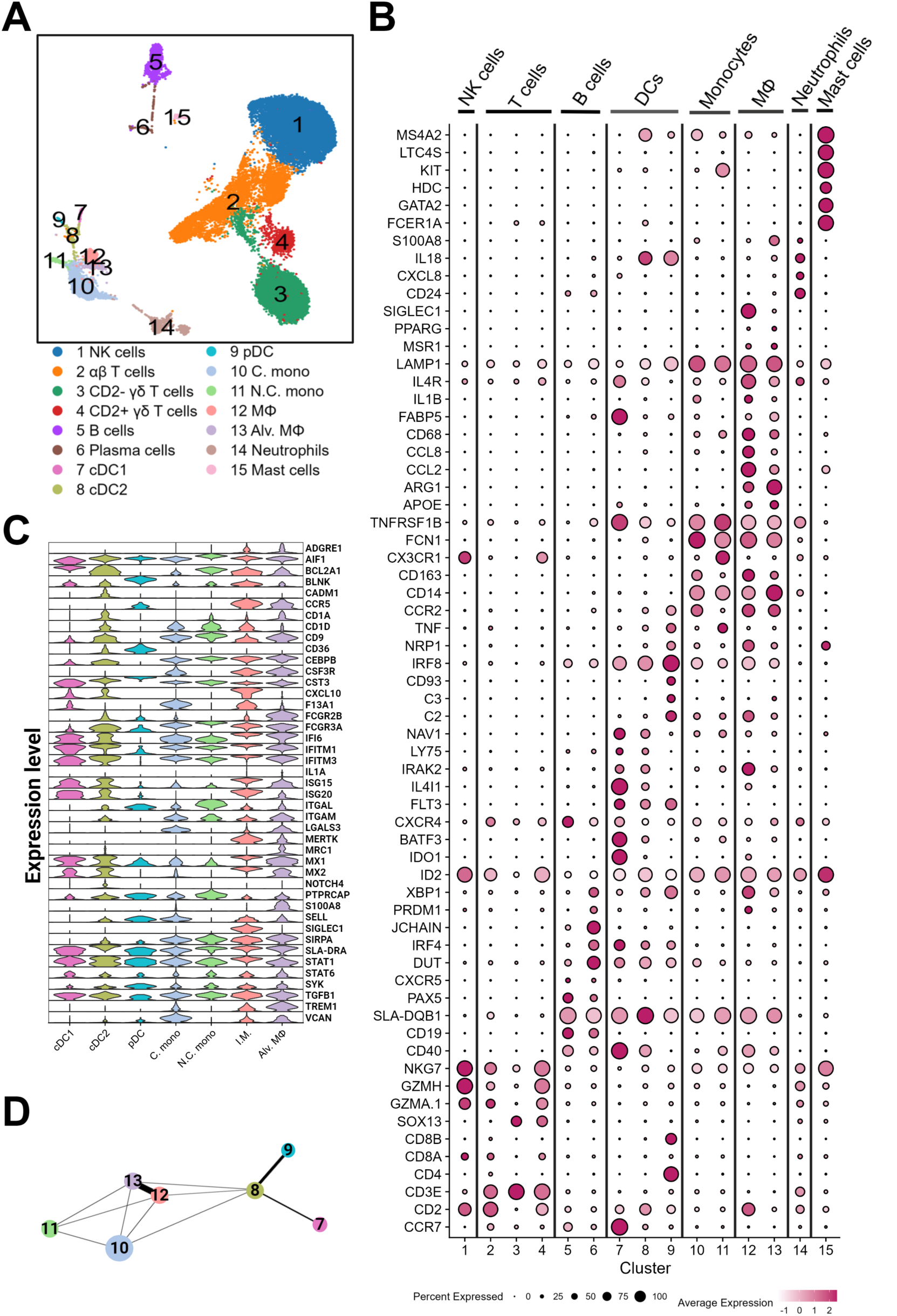
Single-cell transcriptomic analysis of the cellular composition of the pig lung. **(A)** Uniform manifold approximation and projection (UMAP) visualization of lung leukocyte populations colored by cell clusters. Clusters were identified using the graph-based Louvain algorithm at a resolution of 0.5. **(B)** Dot plot showing the Z-scored mean expression of marker genes used to designate cell types to cell clusters. The color intensity indicates the average strength of each marker gene in each cluster. The dot size represents the proportion of cells expressing each marker. Genes with cluster specific increases in expression are presented in Supplementary file 1. **(C)** Expression of marker genes (rows) associated with each myeloid subset. **(D)** A PAGA graph of myeloid states with myeloid cluster nodes colored by cell type.

Most cells were in clusters 1-4, which consisted of closely grouped T and NK cell populations. B cells were separated into a larger cluster (cluster 5) expressing naïve B cell markers (*CCR7*, *CXCR4*, *CD19*^high^) (23,24), and a smaller cluster (cluster 6) expressing plasma cell markers (*IRF4*, *PRDM1*, *XBP1*), antibody secreting (*JCHAIN),* and cell cycling (*DUT*) markers. Partition-based graphical abstraction (PAGA) was applied to infer the trajectory connectivity between myeloid cells (clusters 7-13) (Figure 1D) (25). Dendritic cells (clusters 7-9), monocytes (clusters 10 and 11), and macrophages (clusters 12 and 13) clustered together due to their common enrichment of myeloid cell genes, such as *AIF1*, *CD9*, *CST3*, *SLA-DRA*, and *SIRPA* (CD172α) (Figures 1B and C).

Dendritic cells, identified by their *FLT3* expression, separated into three clusters that correspond to two conventional DC subsets (clusters 7 and 8) and plasmacytoid DC (pDC) (cluster 9). The two conventional DC subsets, designated conventional cDC1 and cDC2 after transcriptionally similar human DC subsets, were both enriched for *CXCL10*, *LY75* (DEC-205), *IRAK2*, *BCL2A1,* and *NAV1* (2,26). Like human cDC1s, cluster 7 cells upregulated the transcription factor *BATF3,* as well as *CXCR4,* and the tolerance induction genes *IDO1* and *IL4I1* (27). cDC2 associated genes in cluster 8 included MHC class II genes, *CADM1*, *CD1A, CD1D*, *FCGR2B*, *ITGAM* (CD11b), *MRC1*, *NOTCH4*, and *SIRPA* (27,28). pDC-associated genes included *CCR2, CD8B*, *IRF8*, *NRP1*, and *TNF*, as well as the B cell receptor signaling genes *BLNK*, *PTPRCAP*, and *SYK* (28–30). Moreover, we similarly detected several genes identified in a previous RNA sequencing analysis of pig peripheral blood pDC (28), including *CD4, CCR5*, *CD36,* the complement system genes *C2*, *C3* and *CD93,* and the transcription factor *XBP1*, which governs the metabolic programing of pDC (31).

Two subsets of monocytes were identified. Cluster 10 was designated as classical monocytes (C. mono; *CCR2*^high^, *CSF3R*^+^, *CX3CR1*^low^, *S100A8*^high^, *SELL*^+^*, VCAN*^+^) while cluster 11 was designated as non-classical monocytes (N.C. mono; *CCR2*^-^, *CX3CR1^high^*, *FCGR3A*^high^, *ITGAL^+^*, *TNF*^+^, *TNFRSF1B^high^*), in agreement with two previously described monocyte subsets in pigs (32–34). Unexpectedly, C. mono expressed more *CD163* transcripts than the N.C. mono subset, which contradicts reports that *CD163* expression is higher on non-classical than classical pig monocytes (32,33).

Macrophages in clusters 12 and 13 expressed multiple macrophage-associated genes (*APOE*, *FABP5*, *LAMP1*, *MSR1*) (34,35). Cluster 12 (MΦ) was enriched for several interferon-stimulated genes (ISGs), including *IFI6*, *IFITM1*, *ISG15*, *ISG20*, *MX1*, *MX2*, and *STAT1* (36–38), chemokines (*CCL2*, *CCL8*, *CXCL10*), and the pro-inflammatory cytokine *IL1B*. The same genes are upregulated by a population of interstitial macrophages found in humanized mice (34). Cluster 12 also upregulated genes associated with M2 type macrophage polarization, including *ARG1* and *IL4R* (39). Cluster 13 (Alv. MΦ) cells expressed genes associated with both immune tolerance (*TGFB1*) and inflammation (*IFITM3*, *IL1A*, *ISG20*). Furthermore, the alveolar macrophage markers *CEBPB*, *LGALS3*, *MRC1* (CD206), and *PPARG* (34,40,41) were enriched in cluster 13 compared to cluster 12.

Two granulocyte populations were detected. Cluster 14 was enriched for neutrophil genes, including *CD24*, *CXCL8*, and *IL18*. Cluster 15 was enriched for mast cell markers, including *FCER1A*, *KIT*, *LTC4S*, and *MS4A2* (42). Consistent with previous publications that performed scRNA-seq on human lung cells (42,43), we found that pig lung mast cells express *GATA2*, which promotes gene transcription to respond to antigenic stimulation (44), and *HDC*, an enzyme related to histamine synthesis (45).

### Characterization of T and NK cells

Next, we performed a detailed analysis of the lung T and NK cell compartments. Clusters 1-4 in Figure 1A were re-clustered at a resolution of 0.7 using the Louvain algorithm, which produced 12 clusters (clusters 1-12). These were annotated according to canonical markers that distinguish mouse and/or human T cell and innate lymphoid cell subsets (Figure 2A and 2B). Trajectory analysis was performed using PHATE, which preserves both local and global relationships between cell clusters (Figure 2C) (46).

**Figure 2.**
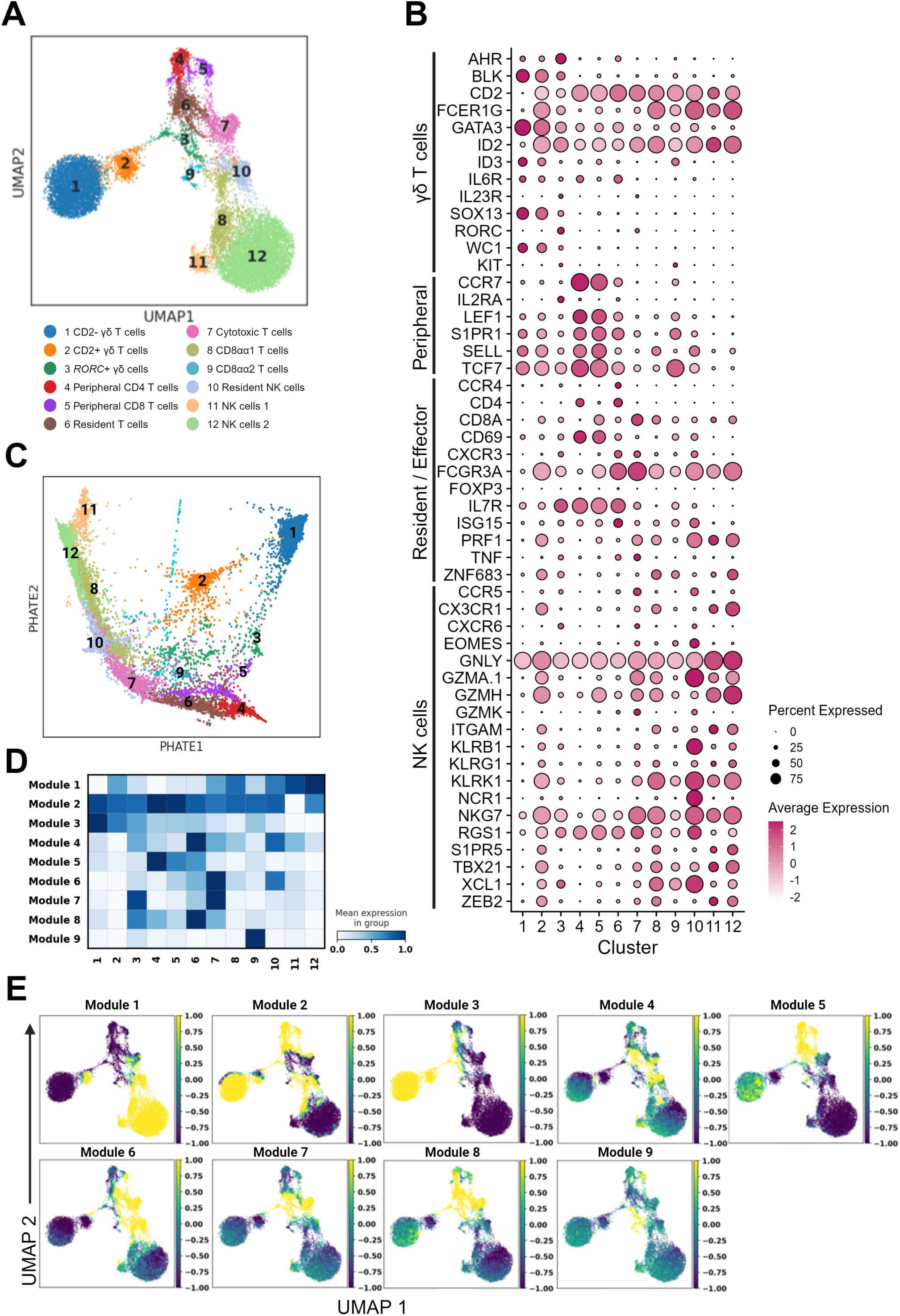
Characterization of T and innate lymphoid cells. **(A-B)** Re-clustered T and NK cell populations from Figure 1A (clusters 1-4) at resolution 0.7 visualized by UMAP **(A)**. **(B)** Dot plot showing the Z-scored mean expression of selected marker genes in clusters from (A). **(C)** PHATE dimensionality reduction method used to visualize same dataset from (A). **(D)** Heatmap of 9 gene modules whose genes had a similar expression pattern across cell clusters. **(E)** UMAP plots showing the expression profiles of genes from modules 1-9. See Supplementary file 1 for a complete list of module genes.

The largest proportion of CD3^+^ cells were γδ T cells, which agrees with the fact that pigs are a high γδ T cell species (47). We identified three clusters of γδ T cells (clusters 1-3). Clusters 1 and 2 respectively correspond to the two major subsets of γδ T cells that exist in pigs, defined as *WC1*^+^*GATA3^hi^CD2*^-^ (CD2^-^) and *WC1^lo^GATA3^lo^CD2*^+^ (CD2^+^) cells. The CD2^-^ subset is found in pigs (47) and other γδ T cell high species such as sheep (48) and cattle (49). This population is resident in a wide range of tissues including lymph node, spleen, liver, intestines, and lung (47,50). CD2^+^ γδ T cells are thought to resemble IFN-γ producing Tγδ1 cells in mice since they express similar transcription factors and seem to require TCR stimulation for their induction (51–53). We found that the CD2^+^ subset was enriched for cytotoxicity-associated genes, such as *CD8A*, *GZMH*, *GZMA.1*, *KLRK1*, and *FCGR3A* (CD16). On the other hand, CD2^-^ γδ T cells upregulated *BLK*, *GATA3*, *ID3*, *IL6R*, and *SOX13* which are expressed by Tγδ17 cells, another major lineage of mouse γδ T cells (54). Prior research has shown that the CD2^-^ population preferentially accumulates in the blood while CD2^+^ subset predominates in lymphoid organs and spleen (55–57). We observed that CD2^-^ γδ T cells were approximately six times more numerous than CD2^+^ cells in the lung. The third γδ T cell subset (cluster 3), which expressed *CD2*, was also enriched in Tγδ17-associated genes (*AHR*, *ID2*, *IL7R*, *IL23R*, *KIT*). However, unlike the CD2^-^ subset, cluster 3 γδ T cells upregulated the transcription factor *RORC* that is required for Tγδ17 lineage commitment. Thus, we designated cluster 3 *RORC*^+^ γδ T cells.

To determine whether any of the pig lung γδ T cell subsets resemble γδ T cell subsets in mice, we subclustered all γδ T cells and integrated them with a scRNA-seq dataset of γδ T cells purified from eight mouse organs (58) (Supplementary figure 2). Mouse γδ T cells separate into eight different subpopulations distinguishable by their expression of *Sell*, *Ly6c2*, *Cd160*, *Gzmb*, *Rorc*, *Areg*, *Klrg1*, and cell cycling genes. Pig *RORC*^+^ γδ T cells separated into two fractions, with the majority overlapping with murine *Rorc*^+^ γδ T cells, the most common γδ T cells in mouse lungs and skin. There was no overlap between CD2^+^ or CD2^-^ γδT cells and any of the mouse γδ T cell subsets, indicating important species-specific differences in the transcriptional landscape underlying γδ T cell subsets in pigs and mice.

Clusters 4-9 represent different subpopulations of αβ T cells. We respectively designated clusters 4 and 5 peripheral CD4 and peripheral CD8 T cells as they displayed a gene profile consistent with naïve peripheral T cells; high *CCR7*, *LEF1*, and *TCF7* (TCF-1) expression and an absence of *IL2RA* (CD25) (59). Cells in cluster 6 upregulated *CD4* and harbored most of the *FOXP3^+^* T regulatory cells (Tregs) and displayed cells of an effector/memory phenotype evidenced by their co-expression of several memory related markers (*CD69, ISG15, IL7R, TNF*). We designated this cluster *CD4* tissue resident T cells due to the enrichment of *CCR4* and *CXCR3,* which are markers of tissue infiltration and residency (60,61). Cluster 7 upregulated *CD8A* and genes associated with cytotoxic effector functions, such as granzyme genes (*GZMH*, *GZMA.1*, *GZMK*), *FCGR3A*, *PRF1* (perforin), and *NKG7* (natural killer cell granule protein 7) and was therefore designated as cytotoxic T cells.

Clusters 8 and 9, respectively designated CD8αα1 and CD8αα2 T cells, resemble T cell populations with a high *CD8A* to *CD8B* ratio that are present in previously published pig thymus and peripheral blood scRNA-seq datasets (62,63). These cells are enriched for *TBX21* (T-bet), *XCL1,* and the transcription factor *ZNF683* (Hobit), which regulates the transcriptional program of several types of tissue resident T cells (64). Compared to CD8αα2 T cells, the CD8αα1 subset upregulated *SELL,* several NK cell-associated genes (*GZMA.1*, *GZMH*, *KLRK1*, *NKG7*) as well as *ZEB2*, a transcriptional repressor that promotes terminal differentiation of CD8^+^ effector and memory T cells (65). Neither CD8αα cluster expressed *NCR1* (NKp46), which identifies a T cell population with a mixed T cell/NK cell phenotype that is prevalent in pig lungs, based on flow cytometry (66). It is notable that CD2^+^ γδ T cells and the CD8αα T cell clusters upregulate many of the same genes, indicating a possible overlap in their functions.

Clusters 10-12 were NK cell populations. Compared to clusters 11 and 12, cluster 10 cells upregulated *EOMES*, a transcription factor expressed by mature NK cells in humans (67), the NK activating receptor *KLRB1,* the chemokine *XCL1,* which encodes a ligand for the XCR1 receptor that is expressed by cross-presenting DC (68), and the transcription factor *TCF7*, which promotes NK cell commitment and survival during maturation (69) (Figure 2B). Furthermore, Cluster 10 had a gene expression profile consistent with human resident NK cells. Compared to other NK cells, cluster 10 cells downregulated *S1PR5* and *KLF2/3* and upregulated *CCR5*, *CD69*, *CXCR3*, *CXCR6*, and *RGS1* (70). Thus, we designated this cluster tissue resident NK cells. Of note, pigs express both NKp46^+^ and NKp46^-^ subsets of NK cells (71). The NKp46^+^ subset has a more activated phenotype and preferentially accumulates in the lung during influenza virus infection (66). Cluster 10 cells may correspond to this NKp46^+^ subset since this population expressed the highest level of *NCR1.* Clusters 11 and 12 both expressed a gene profile consistent with terminally differentiated NK cells from peripheral circulation, including *TBX21* (T-bet) and the leukocyte-adhesion molecule *ITGAM* (67,72,73). They also upregulated *CX3CR1* and *S1PR5,* which are markers of recent lymphatic egress (74,75), the granzyme genes *GZMH* and *GNLY, FCGR3A*, which identifies mature cytotoxic NK cells (76), and *ZNF683,* which is critical for type 1 innate lymphoid cell effector differentiation (77). The main difference between cluster 11 and cluster 12 cells is that the latter upregulated *CX3CR1*, *S1PR5*, *GZMH*, *KLRK1*, and multiple ribosomal protein encoding genes.

To better distinguish functional differences among our various NK and T cell populations, we analyzed transcriptional variation between cells using a Z score matrix, which organizes clusters into modules with similar gene expression patterns (78) (Figure 2D and 2E). This identified 9 modules of co-regulated genes (Supplementary file 1). Module 1, which was enriched in CD8αα T cells, CD2^+^ γδ T cells, and NK cells, was composed of cytotoxic genes, including granzymes (*GZMA.1*, *GZMH*, *GZMM)*, NK cell receptors (*KLRG1, KLRK1*), *TBX21,* and *FCGR3A*. Module 2 was comprised mostly of ribosomal protein encoding genes that varied with peripheral T cells, γδ T cells, resident NK cells, and CD8αα T cells. Module 3, which was enriched in γδ T cells, peripheral T cells, and CD8αα T cells, included many γδ T cell lineage genes (*GATA3*, *RHEX*, *SRGN*, *YBX3*). Module 4, which was upregulated in resident T cells and cytotoxic T cells, included several ISGs and antiviral genes, such as *IFI6*, *IFIT1*, *IRF7*, *ISG15*, *ISG20*, *HERC5*, *MX1*, and *MX2*. Module 5, which varied with peripheral CD4 T cells, resident T cells, and to a lesser extent CD2^-^ and *RORC*^+^ γδ T cells, consisted of genes associated with recently activated T cells, including *CCR7*, *IL7R*, *DAPL1*, *CD40LG*, *CD5, TNFRSF4* (OX40), and *CD69*. Module 6, which was enriched in CD8^+^ T cell subtypes and in resident NK cells, contained several MHC class II molecule-encoding genes, which are expressed by cytotoxic CD8^+^ T cells in pigs. Module 6 also included the granzyme gene *GZMK,* as well as *CXCR3*, which enables certain CD8^+^ T cells to populate the airways (60), and *SH2D1A,* which encodes an adapter protein that regulates signals triggered by SLAM family receptors in T and NK cells (79). Module 7 grouped together *RORC*^+^ γδ T cells, cytotoxic T cells, and a large fraction of resident T cells, due to their common expression of *TNF*, the exhaustion marker *LAG3*, the co-stimulating molecule *JAML,* and the Th17-related genes *IL23R* and *RORC*. Module 8 that contained several Treg markers (*IL2RA*, *CTLA4*, *FOXP3*, *CCR4*, *ICOS*) was enriched in peripheral CD4 T cells, *RORC*^+^ γδ T cells, cytotoxic T cells, and tissue resident T cells. Finally, module 9, which was enriched in peripheral CD8^+^ T cells and a portion of *RORC*^+^ γδ T cells, expressed the cytotoxic lymphocyte-associated genes *CD8B* and *FCRL6*, and general regulators of immune cell physiology, such as *PRDM16*, *CRLF1*, and *NPY*.

### Cross-species comparison of pig, mouse, and human lung leukocytes

To compare the transcriptional profile of pig lung leukocytes with humans and mice, we integrated our dataset with published lung scRNA-seq data from three healthy humans (46-year-old male, 75-year-old male, and a 51-year-old female) (2) and six 10-15-week-old C57BL/6JN mice (3 females and 3 males) (22). After completing the quality control and clustering analyses, we identified 19 clusters (Figure 3A). Although there was good agreement for most lymphoid clusters, the pig dataset contained a substantially lower proportion of myeloid cells (9.75%) compared to the human (57.07%) and mouse (71.25%) datasets (Supplementary file 1). In previous flow cytometry analyses, pig lung myeloid cell populations comprised 68% of live cells (80). The difference with the current study could in part be due to variation in cell isolation protocols which used different enzymes digestion, enrichment, and FACS sorting methods (2,22).

**Figure 3.**
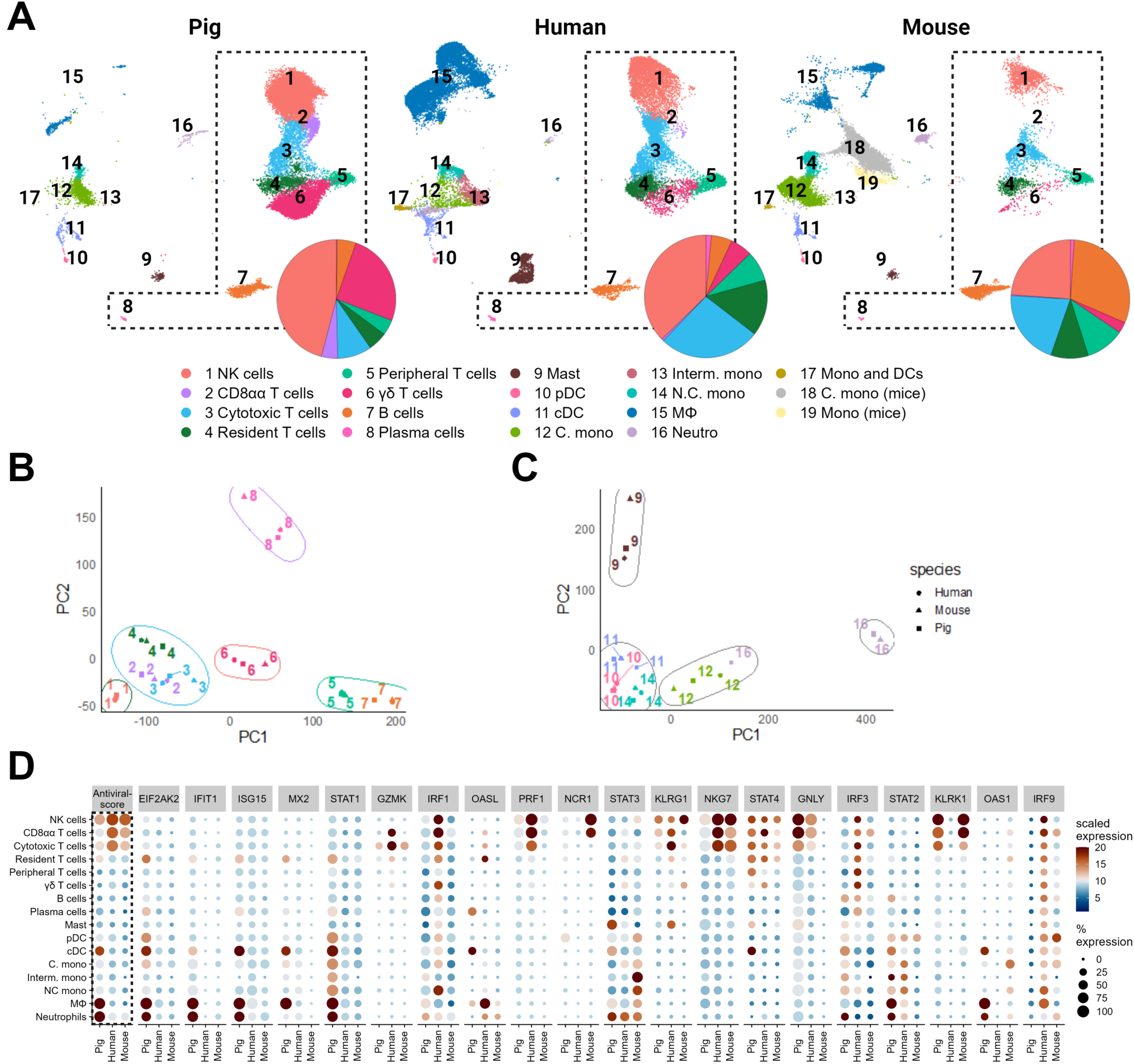
Integrative analysis of pig, human, and mouse lung leukocytes. **(A)** UMAP displaying an integrative analysis of human, mouse, and pig lung leukocytes using a canonical correlation approach to identify shared genes between datasets. Pie charts show differences in lymphoid clusters as a proportion of total lymphocytes. Public datasets containing mouse (22) and human (2) lung samples were used. **(B)** Principal component analysis (PCA) of the transcriptional variation among cell clusters of pigs, humans, and mice. **(C)** Dot plot displaying a cross-species comparison of prototypic antiviral genes and ISGs. Dot size indicates the percentage of each cell expressing the gene. Color saturation represents the strength of expression in positive cells. An antiviral score was calculated by taking the average expression of all 34 genes on a per-cell basis. See Supplementary file 1 for a complete list of genes used.

Among lymphoid cell types, the most notable species differences were that pigs had a substantially greater proportion of γδ and CD8αα T cells compared to mice and humans (Figure 3A). Additionally, pigs and humans had much higher levels of NK cells than mice. In contrast, B cells made up a much greater fraction of mouse lung lymphocytes than in pigs or humans, which agrees with previous scRNA-seq cross-species analyses of lung cells (43,81). Mouse and human lungs harbored a higher proportion of peripheral, resident, and cytotoxic T cells than pigs, perhaps in compensation for their comparative lack of γδ T cells.

Next, we compared cell type-specific clusters across species according to their gene expression patterns using a principal component analysis (PCA) (Figure 3B and 3C). In general, most cell types clustered together across species. The most divergent cell types were plasma cells and mast cells where pigs and humans were more similar to each other than to mice, and neutrophils where human neutrophils were found to be more closely related to human classical monocytes than to mouse or pig neutrophils.

To compare the antiviral machinery in pig, mouse, and human lung immune cells, we analyzed 34 prototypic antiviral genes and ISGs by cell type across species (Supplementary file 1), several of which are plotted in Figure 3D. Examples of cross-species differences include that compared to the other two species (i) porcine DC, macrophages, and neutrophils had higher *EIF2AK2* (Protein kinase R), *IFIT1*, *ISG15*, *MX2*, and *STAT1*, (ii) human cytotoxic cell types expressed higher *GZMK*, *IRF1*, and *PRF1* levels, and (iii) mouse NK and CD8αα T cells expressed more *NCR1*. Additionally, *NKG7* was enriched by human and mouse cytotoxic cell types while *KLRK1* was enriched by pig and mouse cytotoxic cell types. An across-species antiviral score was developed by averaging cell-type expression of all antiviral genes that had orthologues present in all three species. This showed a similar enrichment intensity in NK cells, CD8αα, and cytotoxic T cells, with somewhat higher gene expression in humans. However, pig cDC, macrophages, and neutrophils were the only myeloid cell types among the three species that presented a strong antiviral signature.

### Transcriptional changes induced by influenza infection

We compared lung leukocytes from our mock-infected healthy pigs (Healthy; 19,994 cells, 1,378 genes, 3,728 transcripts) to seven pigs infected with pdmH1N1 influenza virus (FLU; 25,460 cells, 1,669 genes, 4,662 transcripts) and seven pdmH1N1-infected pigs treated with a five-day course of anti-influenza drug oseltamivir phosphate (FLU/OTV; 33,212 cells, 1,583 genes, 4,528 transcripts) (Supplementary file 1). Cells were isolated from lung tissue at 5 days post infection (dpi), which usually coincides with peak lung inflammation in pigs infected with pdmH1N1 (82,83). After integration with Seurat (84), we identified 18 cell clusters (Figure 4A and 4B) using canonical cell markers (Supplementary file 1). Both the FLU and FLU/OTV groups had significantly higher proportions of B cells and cytotoxic, resident, and peripheral T cells, and a lower abundance of NK cells and γδ T cells compared to Healthy pigs (Figure 4B, Supplementary figure 3A). FLU and FLU/OTV pigs had similar cell compositions.

**Figure 4.**
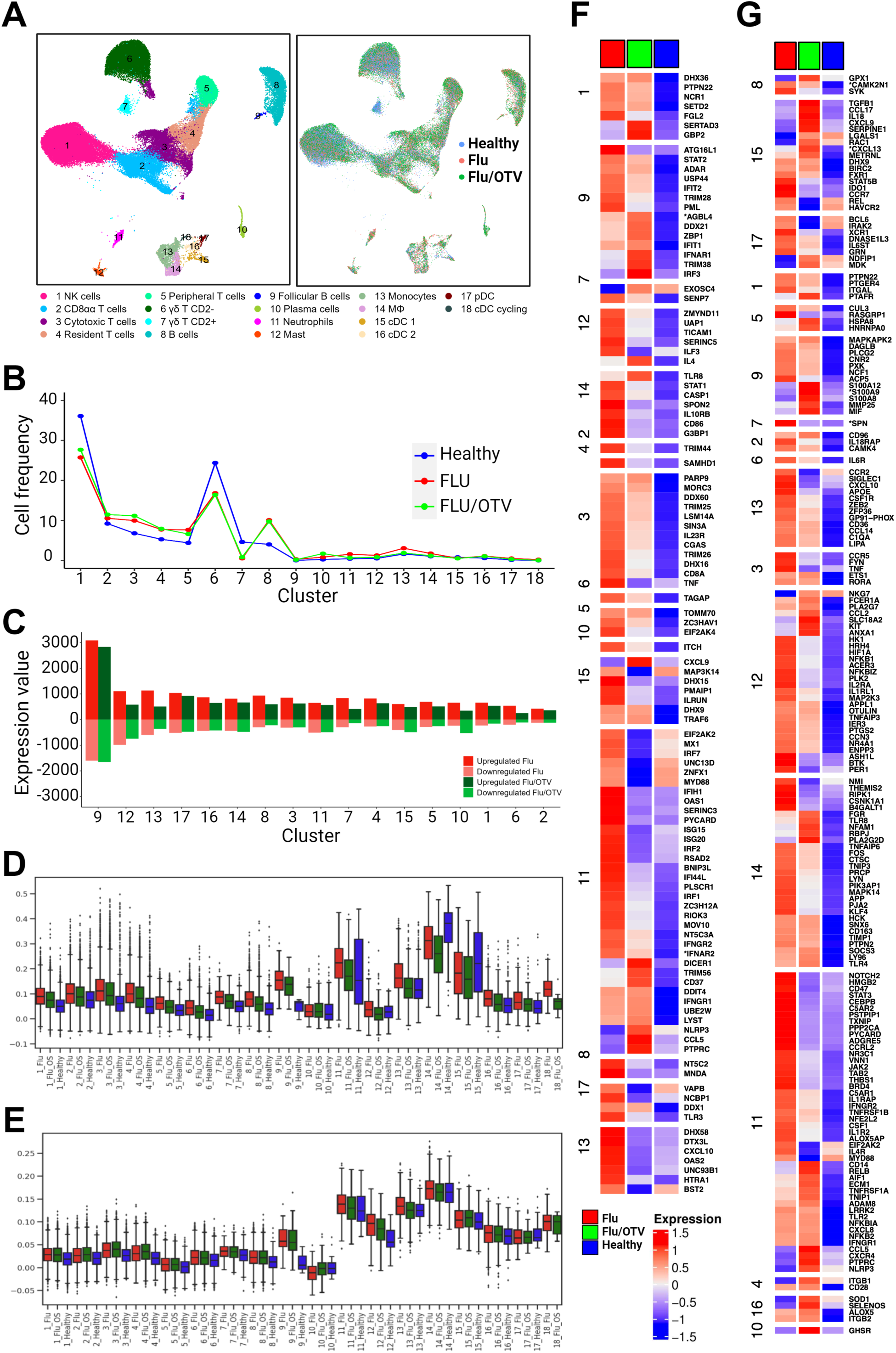
Transcriptional changes induced by influenza infection with and without oseltamivir treatment. **(A)** Visualization of lung leukocytes across different cell types and treatments. UMAPs display an integrative analysis of lung cell types from Healthy, FLU, and FLU/OTV pigs at 5 dpi at resolution 0.5. **(B)** The frequency of each cell type is presented for each treatment. **(C)** Bar graphs displaying the number of upregulated and downregulated differentially expressed genes (DEGs) in FLU and FLU/OTV compared to Healthy pigs. **(D-E)** Box plots showing gene expression score of “Defense Response to Virus” (GO: 0051607) **(D)** and "Inflammatory Response” (GO: 0006954) **(E)** genes for each cell type in their respective treatments. **(E-F)** Heatmaps of the top FLU and FLU/OTV DEGs intersecting the GO terms “Defense Response to Virus” **(F)** and Inflammatory Response” **(G)** for each cell type.

Cells were assessed for the presence of pdmH1N1 influenza viral RNA (Supplementary figure 3B). Only professional antigen-presenting cell types, including B cells, monocytes, macrophages, cDC, and pDC, from FLU and FLU/OTV pigs harbored low viral transcripts. We also compared our lung results to a scRNA-seq dataset generated from the tracheobronchial lymph nodes (TBLN) of FLU and FLU/OTV pigs that were collected at the same time as the lung cells (5 dpi) (Supplementary figure 3C). Compared to the lung samples, TBLNs had fewer NK cells, cytotoxic T cells, and γδ T cells, and higher concentrations of B cells. TBLNs from FLU and FLU/OTV pigs harbored similar proportions of most cell types (Supplementary figure 3D).

Next, we compared DEGs within individual cell types from Figure 4A (Figure 4C and Supplementary file 1). Between Healthy and FLU pigs, the most transcriptionally variable populations were follicular and germinal center B cells (*SELL*^+^*CD22*^+^*CD24*^+^*CD38*^+^ CCR7+*CXCR5^+^CXCR4*^+^) (4,687 genes), followed by mast cells (2,087 genes), monocytes (1,731 genes), pDC (1,557 genes), cDC 2 (1,307 genes), and macrophages (1,253 genes). Between Healthy and FLU/OTV pigs, the most affected cell types were follicular and germinal center B cells (4,486 genes), followed by pDC (1,400 genes), mast cells (1,328 genes), macrophages (1,145 genes), cDC2 (1,075 genes), and neutrophils (1,061 genes).

Since IAV infection causes widespread immune responses in the lung, we calculated the average gene expression scores for genes related to the GO terms “defense response to virus” (GO: 0051607) and “inflammatory response” (GO:0006954) for each cell type (Figures 4D and 4E). Myeloid cell types, including neutrophils, monocytes, macrophages, and DC, had higher scores than lymphoid cell types, largely because they constitutively expressed higher fold change of antiviral and inflammatory genes, even in Healthy pigs. Treatment differences were more pronounced in lymphoid cell types, particularly as regards “defense response to virus” genes. Follicular and germinal center B cells appeared especially reactive to IAV infection, generating a variety of inflammatory and antiviral factors in response (Supplementary file 2). Next, we looked for upregulated DEGs between FLU or FLU/OTV and Healthy pigs that were associated with each GO term (Figure 4F and 4G). Of the 252 genes included in the “defense response to virus”, 105 and 92 were respectively upregulated in FLU and FLU/OTV pigs. Of the 700 genes listed in the “inflammatory response”, 279 and 252 were respectively upregulated in FLU and FLU/OTV pigs (Supplementary file 2). Between FLU and Healthy pigs, neutrophils, follicular and germinal center B cells, and cytotoxic T cells harbored the most “defense response to virus” DEGs, while neutrophils, macrophages, and mast cells contributed the most “inflammatory response” DEGs. Oseltamivir therapy reduced the expression of numerous DEGs upregulated in FLU pigs. However, a small number of DEGs were more upregulated in FLU/OTV than FLU pigs, such as *CXCR4*, *NLRP3*, *TNFRSF1A* in neutrophils, *CXCL9* in DC and *TLR8* and *IL10* in macrophages. This shows that oseltamivir therapy alters how pulmonary immune cells react to IAV infection, possibly because it changes the kinetics of IAV replication in the lung (85).

Next, we performed an Ingenuity Pathway Analysis (IPA) of canonical cellular immune response pathways using DEGs between FLU or FLU/OTV and Healthy pigs to identify IAV-upregulated regulatory networks. For this, we focused on lymphoid cell types which were re-clustered at a resolution of 0.4 (Figure 5A). Pathways upregulated across multiple cell types included “IL-6 Signaling”, “Th2 Pathway”, “Natural Killer Cell Signaling”, “Crosstalk Between Dendritic Cells and Natural Killer Cells”, and “Immunogenic Cell Death Signaling Pathway” (Figure 5B). In response to IAV infection, resident αb T cells, CD2^+^ γδ T cells, and CD8αα T cells upregulated the most pathways (48, 42, and 37 respectively), while peripheral CD4 and CD8 T cells, and innate lymphoid subsets upregulated the fewest (12, 2, and 2 respectively). Pathways such as “Toll-like Receptor Signaling”, “TNFR1 Signaling” and “4-1BB Signaling in T Lymphocytes” were specifically increased by NK cells 2, resident T cells, and resident NK cells. FLU/OTV pigs had fewer upregulated pathways compared to FLU pigs for most cell types suggesting this therapy is having the predicted effect.

**Figure 5.**
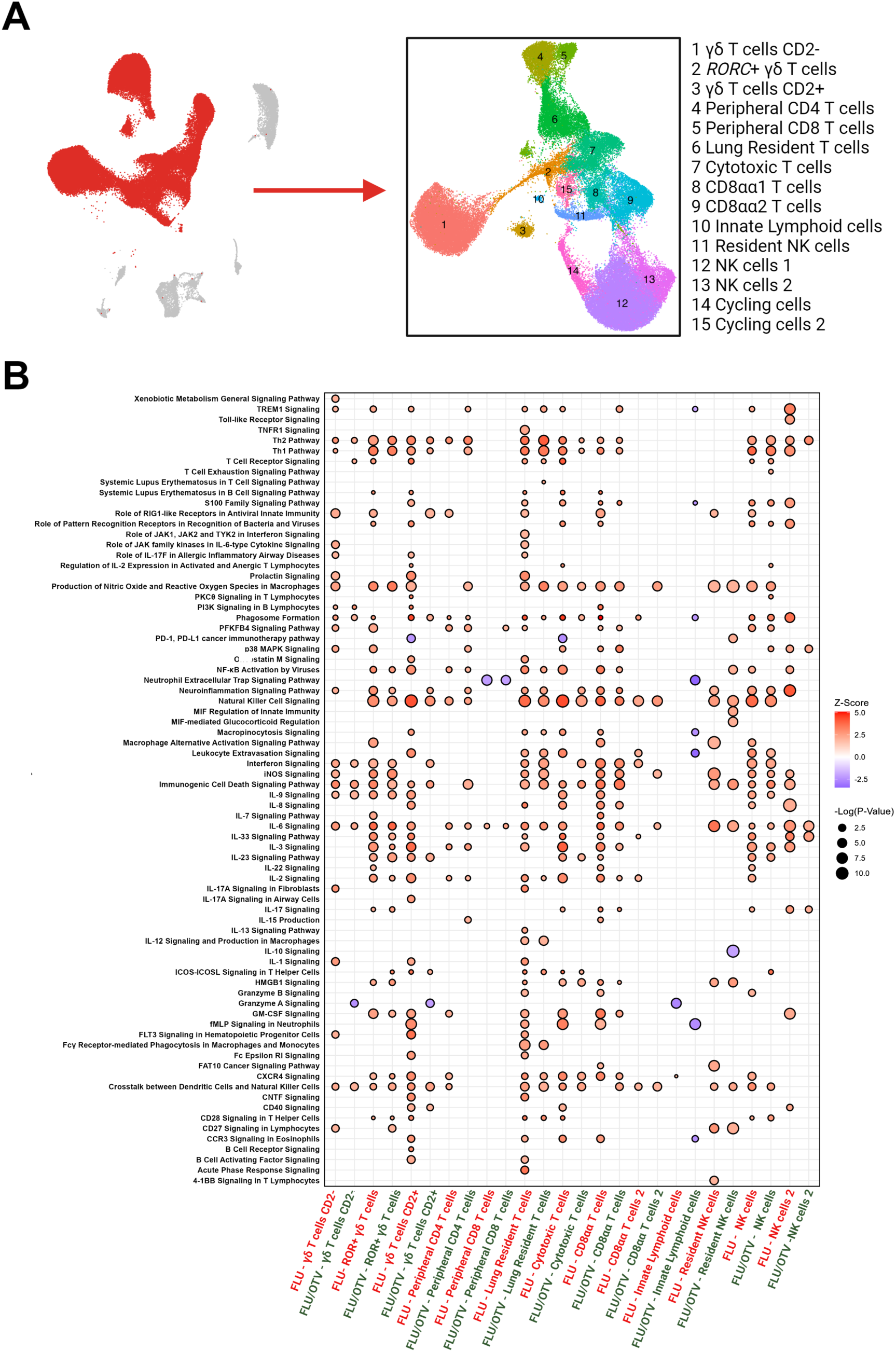
Pathway enrichment analysis of cell types in FLU and FLU/OTV pigs. **(A)** UMAP visualization of T and NK cells reclustered at resolution 0.4. **(B)** Enrichment of canonical cellular immune response pathways for cell types in **(A)** according to ingenuity pathway analysis. The y-axis displays the enriched pathways. A z-score of -2.0 <Z> 2.0 was considered significant. The x-axis displays the cell types in FLU (red font) and FLU/OTV (black font). The dot size displays significance [-log10 (p-value)] of gene sets. Dot color saturation represents the Z score of genes within a network. See Supplementary file 2 for a complete list of genes in each immunological network.

Among the different γδ T cell subsets, FLU CD2^+^ γδ T cells upregulated almost double the number of pathways upregulated by CD2^-^ γδ T cells (42 and 23 respectively). Furthermore, CD2^+^ γδ T cells, in addition to containing most of the CD2^-^ γδ T cell networks, upregulated several cytokine signaling and APC licensing pathways that were absent in CD2^-^ γδ T cells, including “NF-κB Activation by Viruses”, “Natural Killer Cell Signaling”, “Leukocyte Extravasation Signaling”, “IL-17 Signaling in Airway Cells”, “CD40 Signaling”, “B cell Activating Factor Signaling”, as well as multiple cytokine signaling pathways (IL-8, IL-33, IL-3, IL-23, IL-2). It is notable that CD2^+^ γδ T cells upregulated many of the same immune networks that conventional CD4^+^ T-helper cells present during viral infections. Pathways expressed by CD2^-^ γδ T cell that were absent in CD2^+^ γδ T cells included “Role of JAK family kinases in IL-6-type Cytokine Signaling” and “IL-17A Signaling in Fibroblasts”. Interestingly, FLU *RORC*^+^ γδ T cells presented a mixed phenotype, upregulating pathways shared and distinct between CD2^+^ and CD2^-^ γδ T cells. Upregulated pathways unique to FLU *RORC*^+^ γδ T cells included “Macrophage Alternative Activation Signaling Pathway”, “IL-7 Signaling Pathway”, “IL-22 Signaling”, “IL-17 Signaling”, and “HMGB1 Signaling”.

### Effect of influenza infection on cell-cell lung leukocyte communication

We used CellChat (v.1.6.1), a tool for analysis and visualization of cell-cell interactions, to identify putative intercellular communication networks between pulmonary immune cell subsets in FLU or FLU/OTV versus Healthy pigs. CellChat objects were created from an integrated Seurat dataset containing similar numbers of cells from each treatment group (Supplementary figure 4A). Overall, FLU pigs presented a considerable increase in the number and strength of inferred cell-cell interactions compared to Healthy pigs. Many of these interactions were substantially reduced or undetectable in FLU/OTV versus Healthy pigs (Supplementary figures 4B and 5).

We analyzed immune-related receptors and ligands upregulated in FLU versus Healthy pigs within cell type. Resident T cells upregulated the ligands *CD6*, *PASP*, *CLE2B*, *CD48*, and *CD40LG* that respectively bind the receptors *ALCAM*, *GPR37*, *KLRB1*, *CD244*, and *CD40*, which are expressed by multiple cell types (Figure 6A). The most connected receptor in resident T cells was *CCR4* (Figure 6A), which facilitates T cell transmigration from the pulmonary vasculature into the interstitial compartment (86). The CCR4 ligand *CCL5* was upregulated principally by NK cells, CD8αα T cells, cytotoxic T cells, and CD2^-^ γδ T cells. Peripheral T cells upregulated *ICAM2* that was connected to the integrins *ITGAL*_*ITGB2* (LFA-1) and *ITGAM*_*ITGB2* (MAC-1) (Supplementary figure 6A), which mediate leukocyte adhesion. These integrins were upregulated by a wide variety of cell types in response to IAV infection.

**Figure 6.**
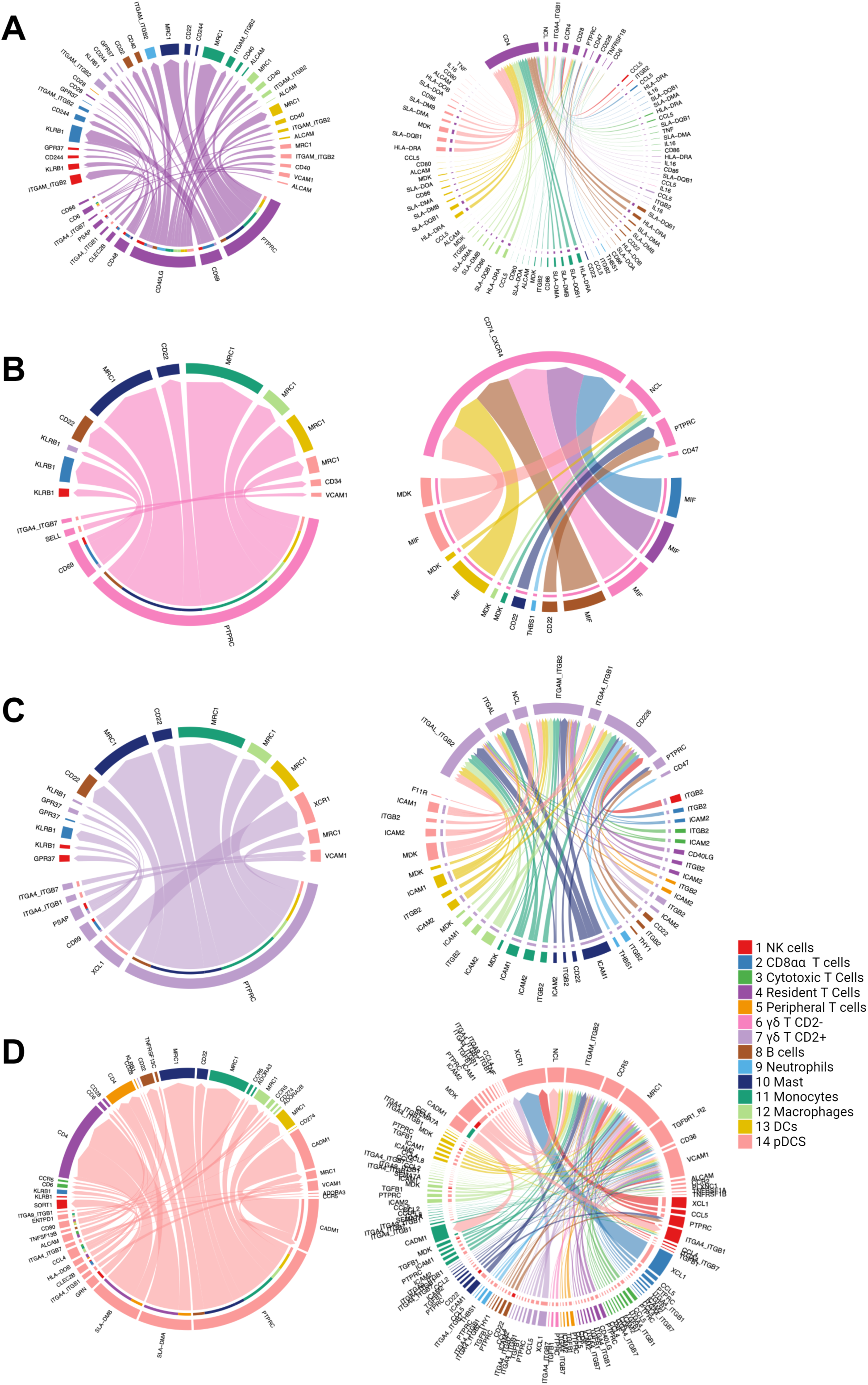
Cell-cell interaction changes associated with influenza infection. **(A-H)** Circle plots showing upregulated ligand (left panel in each pair) and receptor (right panel) of FLU versus Healthy pigs in tissue resident T cells **(A)**, CD2^-^ γδ T cells **(B)**, CD2^+^ γδ T cells **(C)**, and pDC **(D)**. The size of the interaction arrow is in accordance with the transcriptional level of ligand or receptor genes in each cell type.

As regards γδ T cells, only the CD2^-^ subset upregulated *SELL* in response to influenza infection, which was connected via one of its receptors, *CD34*, to pDC (Figure 6B). Like peripheral T cells, CD2^-^ γδ T cells upregulated the heteromeric macrophage migration inhibitory factor (MIF) receptor formed by CD74 and CXCR4 (Figure 6B). MIF is a pleiotropic cytokine secreted by multiple cell types during an influenza infection (87). While MIF helps to restore the alveolar epithelium after injury (88), mouse studies have demonstrated that MIF can also exacerbate influenza-associated pathology by decreasing antiviral type I/III IFN levels (87). The strong *MIF*-*CD74*_*CXCR4* connections in the current dataset suggest a role for this cytokine in swine anti-influenza immune responses. The expression of *XCL1*, which is linked to *XCR1* on pDC, distinguished CD2^+^ from CD2^-^ γδ T cells (Figure 6C). This connection suggests that porcine CD2^+^ γδ T cells help attract *XCR1*-expressing pDC into the lung during influenza virus infections.

Among the myeloid mononuclear cell types, IAV infection caused monocytes to upregulate ligands that increased signaling from MHC class II and T cell costimulatory genes to CD4 expressing cells, and through *ICAM1-ITGB2*, *CCL14-CCR1*, and *CXCL10-CXCR3* ligand-receptor pairs (Supplementary figure 6). Furthermore, monocytes upregulated receptors that enhanced signaling through *TGFB1-TGFbR1_R2*, *RETN-TLR4*, *CCL5-CCR1*, *CCL4-CCR5*, and *MIF-CD74_CXCR4* ligand-receptor pairs (Supplementary figure 6B). Ligands upregulated by macrophages but not monocytes include *CD48* that interacts with *CD244* (2B4) on NK cells and CD8αα T cells, as well as *CXCL2* and *CXCL8* (IL-8) which interact with *CXCR2* on neutrophils (Supplementary figure 6C). CD48-CD244 interactions mediate NK-cell self-tolerance (89) while CXCL2 and CXCL8 are chemotactic for polymorphonuclear leukocytes (90,91). Macrophages were also connected via *TNF* to cytotoxic CD8 T cells and pDC through upregulated *TNFRSF1B* receptor (Supplementary figure 6C).

Influenza infection enhanced cDC and pDC interactions with CD4 T cells through upregulation of *SLA*-*DMA* and *SLA*-*DMB* (Figure 6D and Supplementary figure 6D). Additionally, both DC subtypes connected with *MDK*, *ICAM1*, and *ICAM2* expressing cells through their upregulation of receptors *NCL* and *ITGAM*_*ITGB2*. Unlike pDC, cDC upregulated the ligands *MDK*, *ICAM1*, *PSAP*, *ITGB2*, and *SEMA7A* that respectively connected them to *NCL*, *ITGAM*_*ITGB2*, *GPR37*, *ICAM1*/*ICAM2*, and *PLXNC1* expressing receptor cells (Supplementary Figure 6D). Although usually associated with synapse formation in the olfactory system (92), *SEMA7A* and its receptor *PLXC1* also regulate leukocyte transmigration and monocyte activation during inflammation (93–95). Ligand-based interactions upregulated by IAV in pDC but not cDC include *CCL4* that interacts with *CCR5* on monocytes and macrophages, *CLEC2B* that interacts with *KLRB1* on NK cells and CD8αα T cells, and *GRN* (granulin) that interacts with *SORT1* (sortilin) on NK cells (Figure 6D). Granulin is an essential cofactor for TLR9 in pDC (96). pDC upregulated receptors that enhanced *XCL1*-*XCR1* interactions with NK cells, CD8αα T cells, and CD2^+^ γδ T cells. They also upregulated *TGFbR1*_*R2* and *VCAM1* that respectively connected with *TGFB1* and *ITGA4*_*ITGB1* (VLA-4) in multiple cell types (Figure 6D).

Overall, these results demonstrate a complex interaction network among cells of the innate and adaptive immune systems 5 days after IAV infection and that oseltamivir treatment decreases these interactions.

## DISCUSSION

The current work demonstrates the biological resolution gained from single cell transcriptomic analyses of the pulmonary immune cell types in healthy and influenza-infected five-week-old pigs, that importantly corresponds to human toddlers of approximately five years of age (97). Apart from CD8αα and γδ T cells that were prominent only in pigs, most pig immune cell types overlapped with published mouse and human lung datasets (2,22). Nonetheless, we identified interspecies differences in gene expression within cell types, particularly in the myeloid compartment, including that pig neutrophils, DC, and macrophages constitutively expressed higher levels of several antiviral genes than their mouse and human counterparts, which may have implications for translating certain aspects of pulmonary immunity from pigs to humans.

Our study examined the transcriptional profiles of T and NK cell clusters, which contained both tissue resident and peripheral T and NK cell populations. CD8αα T cells, which were prominent in our pig lung samples, are barely detectable in previously published mouse or human lung datasets (2,22). These cells, and particularly the CD8αα1 subset, resemble a population of NK cell-like T lymphocytes that we previously described in a single cell analysis of pig thymopoiesis, which upregulate T cell memory/activation markers, NK cell signature genes, and the transcription factor *ZNF683* (62). Porcine CD8αα T cells may correspond to a formerly described population of T lymphocytes with a mixed NK and T cell phenotype that are prevalent in the liver and lung of pigs (66). These cells have the capacity to produce perforin and degranulate upon triggering of CD3 or NK receptors. They also proliferate in pig lungs after IAV infection, indicating that they may be involved in controlling virus replication (98).

We identified three transcriptionally distinct populations of γδ T cells, the largest of which corresponds to a previously described pig CD2^-^ γδ T cell subset (55,56,62). Although abundant in pigs, current understanding of CD2^-^ γδ T cells is limited owing to their absence in mice and humans. When compared to a previously published scRNA-seq dataset from pig intestinal ilium (50), another mucosal site, we observed a much higher ratio of CD2^-^ to CD2^+^ cells in lung γδ T cells, indicating that the CD2^-^ subset is enriched in the respiratory compared to the intestinal mucosa.

Lung CD2^+^ γδ T cells had a similar transcriptomic profile to CD2^+^ γδ T cells in pig blood and ileal datasets (50,63), including enrichment of *FCER1G* and the chemokine *XCL1*. However, tissue-specific differences were also found. For instance, lung CD2^+^ γδ T cells were enriched for the pro-inflammatory transcription factor *TBX21*, and granzymes *GZMH* and *GNLY*, which were not detected in ileal CD2^+^ γδ T cells. This points to the acquisition of tissue-specific adaptations, perhaps for different pathogens. The third γδ T cell cluster, *RORC^+^* γδ T cells, expressed a mixture of CD2^+^ and CD2^-^ subset genes. This population was unusual for its high expression of *AHR*, which encodes aryl hydrocarbon receptor (AhR). AhR is a ligand-activated transcription factor that integrates metabolic, microbial, and environmental signals to modulate transcriptional programs in a ligand and cell-type specific context (99). Dendritic epidermal T cells (DETC), a type of γδ T cell that account for nearly all epidermal lymphocytes, express high levels of AhR and play a critical role in tumor surveillance and wound healing (100). Furthermore, a subset of *RORC*-expressing DETC that upregulate *IL7R* are a major source of IL-17A, which is required for wound healing after skin injury (100). Given their enrichment of *AHR*, *IL7R,* and Th17 lineage genes, lung *RORC^+^*γδ T cells may perform similar functions to IL-17A producing DETC within the respiratory tract.

Influenza virus infection and oseltamivir treatment altered the cellular composition, transcriptional networks, and cell-cell interactions in the lung. Notable observations were that IAV infection increased the proportion of T and B cells among lung lymphocytes, while NK cells decreased, which agrees with previous mouse studies that examined anti-influenza immune responses in the lung (101–103). Infection caused a general increase in immune-related DEGs across lymphoid and myeloid cell types. A minor population of B lymphocytes presenting a follicular/germinal center B cell phenotype was the most affected cluster, upregulating the transcription repressors *TRIM28* and *ADAR1*, and type I interferon response genes (*IFR3*, *IFIT2*, *STAT2*) (104–106). Accordingly, lung germinal center B cells may interact with T cells and produce high affinity antibodies in response to influenza virus infection (107). Upon examination of T and NK cell subsets by signaling pathway enrichment tests, CD8αα T cells emerged as particularly reactive to IAV infection, altering a wider range of immune networks than peripheral or resident conventional T cell subtypes, notably cytokine and acute phase response signaling pathways. This ability to deploy diverse effector responses soon after infection is typical of innate-like T cells which specialize in quickly sensing their local environment and transmitting those signals to downstream innate and adaptive effector cells to elicit antimicrobial protection (108).

Our objective was to compare the responses of various subsets of γδ T cells to IAV infection, as these cells comprise a significant fraction of the total T lymphocytes in the lungs of pigs (47,109–111). Furthermore, our examination of γδ T cells in the lungs of pigs unveiled significant interspecies variations, which has ramifications for the transfer of paradigms about the roles of γδ T cells from mice to pigs. CD2^+^ γδ T cells exhibited more reactivity than CD2^-^ γδ T cells, which is consistent with a prior finding that cytokines were generated exclusively by CD2^+^ γδ T cells in pigs infected with IAV (98). A considerable proportion of the pathways that were upregulated in CD2^+^ γδ T cells also exhibited overlap with the pathways induced by IAVs in resident T cells and CD8αα T cells. This finding implies that these three cell types utilize comparable mechanisms to augment the magnitude and caliber of IAV-driven immune responses. It proved challenging to differentiate the specific function of CD2^-^ γδ T cells in comparison to the other subsets, on the basis of IAV-induced immunological networks and cell-cell interactions. Nevertheless, despite their comparatively restricted array of effector functions, the presence of CD2^-^ γδ T cells as the most numerous subtype of single T cells in the pig’s lung and their response to IAV infection indicate that these cells play a substantial part in the pig’s anti-IAV defenses. T cells expressing *RORC^+^* increased a subset of pathways that were enriched in CD2^+^ γδ T cells. The immune networks that were specifically upregulated in *RORC^+^* γδ T cells consisted of IL-7, IL-17, and IL-22 signaling. This finding aligns with the notion that IL-7 modulates IL-22 production by augmenting RORγt expression (112). Furthermore, *AHR*, a protein that is extensively expressed in *RORC^+^* γδ T cells, can directly stimulate the production of IL-22 and the differentiation of Th17 cell subtypes, including Tγδ 17 and DETC. Analogous *RORC^+^* γδ T cells are prevalent in the lungs of mice and offer early immunity against influenza virus infection via a CD1d-dependent mechanism (109).

Treating IAV-infected pigs with oseltamivir substantially reduced viral loads and lung pathology compared to untreated infected pigs (85). This was reflected in the various immunological signaling pathways and cell-cell interactions that were downregulated or undetectable in FLU/OTV pigs compared to FLU pigs across a range of cell types. This was particularly noticeable in cell types known to respond vigorously to virus exposure, such as CD8αα T cells, cytotoxic T cells, CD2^+^ γδ T cells, and NK cells. These data demonstrate that oseltamivir treatment significantly decreased immune activity in the lungs of IAV-infected pigs, which supports that, when given early enough after infection, this medication is effective at reducing IAV-induced clinical signs and inflammation (113,114).

In summary, we constructed a single cell atlas of the porcine pulmonary immune system comprising of immune cells isolated from the lung tissue of healthy pigs, influenza virus infected pigs, and influenza infected pigs treated with oseltamivir. Our results indicate that at 5 dpi, pig lungs are undergoing dynamic changes in cell recruitment, cellular activation, and tissue remodeling in response to ongoing virus replication, and that some of these changes are reduced with oseltamivir administration. Since pigs are increasingly used to model human microbial infections and respiratory disorders, such knowledge is important to determine where pigs may succeed and fail to predict human pulmonary immune responses. Furthermore, our atlas helps understand the porcine anti-influenza defense system, which is important to make progress in the design of more protective IAV vaccines and antiviral therapies. It may also be useful for elucidating several important yet poorly understood aspects of swine IAV immunity such as host factors that influence virus transmission and early life imprinting of the influenza virus-specific memory lymphocyte compartment.

### Limitations of the study

Although this study has provided a comprehensive transcriptional analysis of pig lung leukocytes, we only profiled pigs at a single age and did not include non-immune cell populations. Moreover, lung samples were obtained at a single time point after influenza virus infection. In the future, it will be important to expand our dataset to capture the full extent of lung cellularity at different ages and different timepoints after influenza infection. Additionally, we cannot exclude that some of the interspecies differences we observed in cell subsets and intercellular communication networks were due to biological and technical effects, such as differences in physiological age, tissue preparation methods, and sequencing saturation. Hence, validation of our findings is required using additional datasets as they become available.

## MATERIALS AND METHODS

### Pigs

Cells analyzed in the current work were isolated from a previously published swine influenza challenge study (85). Briefly, fourteen four-week-old commercial mixed breed pigs seronegative for antibodies against H1N1, H3N2, and B influenza viruses were intratracheally inoculated with 1 x 10^6^ TCID_50_ 2009 pandemic H1N1 A/California/04/2009 influenza virus, as previously described (115). Seven of these pigs were orally administered 75 mg oseltamivir phosphate (Lupin Pharmaceuticals) twice a day for five days after infection (FLU/OTV group) while the remaining seven pigs were left untreated (FLU group). Tissue samples were obtained from two additional pigs (Healthy group) that were mock infected with virus-free Dulbecco’s Modified Eagle Medium. At 5 dpi, necropsies were performed as previously described (115), during which lung and lymph node samples were collected for scRNA-seq.

### Tissue sampling and cell isolation

Approximately 1 g of tissue collected from the left cranial, middle, and caudal lung lobes were combined and digested with 2.5 mg/mL of Liberase TL (Roche, Indianapolis, IN) in Dulbecco’s Modified Eagle Medium (Thermo Fisher, Waltham, MA) at 37°C for 45 minutes, passed through a 100 μm cell strainer (Thermo Fisher, Waltham, MA), and treated with an ammonium chloride-based red blood cell lysis buffer. The resulting cell suspensions were washed, stained with propidium iodide for exclusion of dead cells, and FACS sorted for live cells using a Sony SH800 Cell Sorter (Sony Biotechnology, Japan). Ten lung samples were sequenced from pooled cell suspensions of two piglets each, while other six lung samples were individually sequenced (Supplementary file 1). A total of 12 datasets from pooled and non-pooled samples were obtained - 2 Healthy, 5 FLU, and 5 FLU/OTV.

### Single-cell RNA sequencing

Single cell libraries were prepared using the 10x Genomic Chromium Next GEM Single Cell 3’ reagent kit (v3.1) according to the manufacturer’s instructions. Sequencing was performed on an S4 flow cell of the NovaSeq 6000 sequencer (Illumina) to obtain paired end reads.

### Data processing and clustering analysis

Sequence reads were aligned to the pig reference dataset Sscrofa 11.1, followed by creation of barcode gene matrices using Cell Ranger v7.1.0 (10x Genomics). Clustering analyses were performed in R (v4.0.2) using Seurat (v4.3.0.1) (84). Pre-analysis quality control was performed by removing genes expressed in <3 cells and excluding cells with aberrantly high (>5000) or low (<550) gene counts and high mitochondrial gene expression (>9%) (Supplementary file 1). Highly variable genes were used to produce the principal component analysis (PCA). Canonical cell cycle markers were then used to regress out cell cycle effects before dimensionality reduction. Dimensionality reduction was performed using uniform manifold approximation and projection (UMAP). Cell lineages were manually annotated based on algorithmically defined marker gene expression for each cluster. Differentially expressed genes (DEGs) were identified within each cluster using the *FindAllMarkers* function with a minimum Log2 fold change threshold of + 0.25 using a Wilcoxon Rank-Sum test.

### Trajectory inference analysis

We applied partition-based graph abstraction (PAGA) for trajectory analysis using *scanpy.tl.paga* function from Scanpy (v1.9.1) (116), which reconciles clustering and pseudotemporal ordering algorithms and allows the inference of complex cell trajectories and differentiation trees (116). Potential of heat diffusion for affinity-based trajectory embedding (PHATE) (v1.0.10) was also applied to align cells into a developmental trajectory (46). The normalized datasets were imported into PHATE to instantiate a PHATE estimator object with default parameters, and then PHATE embedding was generated within a low dimension, that recapitulated the expected lineage relationships between the clusters, suggesting a progressive differentiation of cell populations.

### Highly correlated gene module identification

We used Hotspot package (v1.1.) to identify highly correlated genes into modules, which computes gene modules by finding informative genes with high local autocorrelation and clustering the results in a gene-gene affinity matrix (78). Briefly, we used the ‘normal’ model on the log-normalized counts to create a hotspot object and construct the K-nearest-neighbors (KNN) graph with 30 neighbors, and then selected the top 500 genes with the highest autocorrelation Z-scores. We then computed pairwise local autocorrelations between these genes, and clustered genes into modules using *create_modules* function (minimum gene threshold of 20, FDR threshold of 0.05, core_only=True). Finally, aggregated gene module scores were calculated using function *calculate_module_scores*.

### Cross-species integration

We used Seurat (v4.3.0.1) to perform cross-species comparisons with published datasets of human lung cells (EGAS00001004344) (2), mouse lung cells (GSE109774) (22) and healthy pigs, and to integrate the influenza infected pig lung and lymph node data. Healthy pig lung γδ T cells were integrated with a dataset of mice γδ T cells (58). The Ensembl genome browser (Ensembl Genes 105) was used to convert human (GRCh38) and mouse (GRCm39) gene names to the corresponding pig names prior to integration (https://www.ensembl.org/biomart/martview/). Only genes with one-to-one orthologs were included in the analyses. The datasets were similarly transformed using common cut-offs for low quality genes and cells as described above. Each dataset was independently normalized before identifying the most variable features, after which a standard integration workflow was followed as previously described (62). Briefly, the *FindIntegrationAnchors* function identified a set of anchors (pairs of cells from each dataset that are contained within each other’s neighborhoods) between datasets using the top 30 dimensions from the canonical correlation analysis to specify the neighbor search space. Next, an integrated dataset was created by running the *IntegrateData* function. Then, the clustering analysis was performed as above. PCA analysis was performed based on the average gene expression profile of highly variable genes across cell types within different species.

### Differentially expressed gene analysis within cell types between FLU, FLU/OTV, and Healthy pigs

Within each cell type, we identified the differently expressed genes (DEGs) between FLU and Healthy pigs, and between FLU/OTV and Healthy pigs using Seurat *FindMarkers* in default settings, which performs differential expression testing based on the non-parametric Wilcoxon rank sum test. DEGs were examined for enrichment in the Gene Ontogeny Database (http://geneontology.org) using the terms of “inflammatory response" (GO:0006954) and “defense response to virus” (GO:0051607) and the R package clusterProfiler. DEGs were subjected to Ingenuity Pathway Analysis (IPA, Qiagen) to identify immune-related canonical pathways and gene networks that are most activated between the FLU or FLU/OTV and Healthy pigs. A z-score of -2.0 <Z> 2.0 was considered significant (117).

### Cell signaling gene scoring

The *scanpy.tl.score_genes* function was used to quantify the gene expressions of antiviral [“defense response to virus” (GO:0051607)] and inflammatory pathways [“inflammatory response" (GO:0006954)] in each cell. The signature score for each cell was defined as the average expression of selected genes subtracted with the average expression of reference genes. The reference gene set was randomly sampled from the gene pool for each binned expression value.

### Cell-cell interaction analysis

Cell-cell interactions were analyzed with the CellChat (118) package (v1.6.1), using the standard pipeline. We converted the published human dataset of ligand receptor pairs (118) to pig gene names, which then was used as a reference to visualize upregulated ligand-receptor interactions within different cell types in FLU or FLU/OTV versus Healthy pigs. Interaction circular maps were built using the literature supported ligand-receptor pairs and connecting the edges between them using the CellChat function *netVisual_chord_gene*. The size of the interaction arrow is in accordance with the transcriptional level of ligand or receptor genes in each cell type.

### Data availability

The sequencing data are available at Gene Expression Omnibus (accession GSE257548). The final data are available for download and direct query at https://singlecell.broadinstitute.org/single_cell/study/SCP2550/single-cell-sequencing-of-influenza-infected-pig-lungs-and-lymph-node; https://singlecell.broadinstitute.org/single_cell/study/SCP2551/transcriptional-changes-induced-by-influenza-infection-in-pig-lungs; and https://singlecell.broadinstitute.org/single_cell/study/SCP2552/characterization-of-t-and-nk-cells-in-pig-lungs. R and Python scripts used for processing of scRNA-seq data are available at: https://github.com/Driver-lab1/sc_RNAseq_Pig_Lung_2024. Any additional information required to reanalyze the data reported in this paper is available from the lead contact upon request.

## Supporting information

Suppementary Figures

## Acknowledgments

We would like to thank the University of Missouri Bioinformatics and Analytics Core for their assistance with high performance computing data management and data transfer.

## Supplementary figure legends

**Supplementary figure 1. scRNA-seq data statistics.** Box plot of **(A)** mean number of cells, **(B)** gene distribution per cells (log_10_), **(C)** transcript distribution per cell (log_10_), and (**D)** UMIs detected per cell (log_10_) for each dataset. Public datasets containing mouse (22) and human (2) lung samples were used.

**Supplementary figure 2. Integration of pig and mouse** γδ **T cells.** UMAP plots showing **(A)** merged dataset containing pig and mouse cells, **(B)** pig and mouse γδ T cells by tissue of origin, original mouse γδ T cell type annotation, and **(D)** pig γδ T cell type annotation in Figure 2A.

**Supplementary figure 3. Transcriptional changes induced by influenza infection and oseltamivir treatment. (A)** UMAP displaying an integrative analysis of lung cells from Healthy, FLU, and FLU/OTV pigs at 5 dpi. **(B)** Dot plot displaying a cross-treatment comparison of influenza genes. Dot size indicates the percentage of each cell expressing the gene. Color saturation represents the strength of expression in positive cells. Grey dots indicate absence of influenza gene expression. **(C)** UMAP displaying an integrative analysis of lung and TBLN cells from combined FLU and FLU/OTV pigs at 5 dpi. Matched lung and TBLN samples were collected from three pigs in each treatment. Pie charts show differences in leukocyte clusters as a proportion of total cells. **(D)** Bar graphs displaying the frequency of each cell type in TBLN for each treatment. **(E)** UMAP plots showing gene expression score of “Defense Response to Virus” (GO: 0051607) and "Inflammatory Response” (GO: 0006954) in pig lung leukocytes.

**Supplementary figure 4. Influenza and oseltamivir treatment induces changes in cell-cell communication. (A)** UMAP visualization of cell clusters used in cell-cell communication analysis. B cells and plasma cell clusters from Figure 4A were grouped together as were cDC and proliferating cDC cells. **(B)** Heatmaps showing the relative number and strength of predicted cell communication pathways for each cell type in FLU (upper) and FLU/OTV (lower) compared to Healthy. The Y-axis represents outgoing signaling, and X-axis represents incoming signaling from each cell cluster.

**Supplementary figure 5. Cell-cell interactions upregulated in influenza infected pigs treated with oseltamivir by cell type. (A-H)** Circle plots showing upregulated ligand (left panel in each pair) and receptor (right panel) of FLU/OTV versus Healthy pigs in **(A)** resident T cells, **(B)** peripheral T cells, **(C)** CD2^-^ γδ T cells, **(D)** CD2^+^ γδ T cells, **(E)** monocytes, **(F)** macrophages, **(G)** cDC, and **(H)** pDC. The size of the interaction arrow is in accordance with the transcriptional level of ligand or receptor genes in each cell type.

**Supplementary figure 6. Additional cell-cell interaction changes associated with influenza infection by cell type. (A-D)** Circle plots showing upregulated ligand (left panel in each pair) and receptor (right panel) of FLU versus Healthy pigs in **(A)** peripheral T cells, **(B)** monocytes, **(C)** macrophages, and **(D)** cDC. The size of the interaction arrow is in accordance with the transcriptional level of ligand or receptor genes in each cell type.

