## Supplementary material for "Single cell sequencing of pig lungs reveals immune responses underlying influenza infection and oseltamivir therapy": Suppementary Figures

### Supplementary figure 1

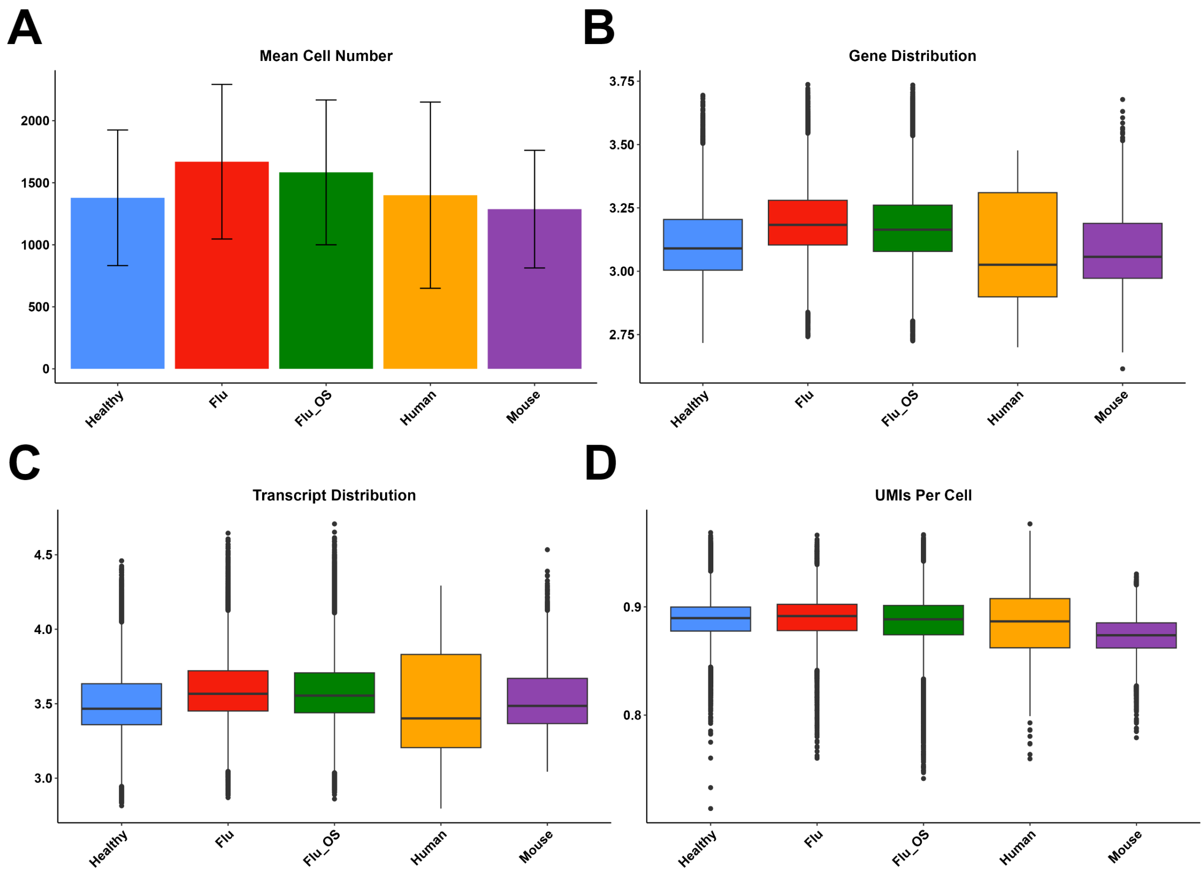

### Supplementary figure 2

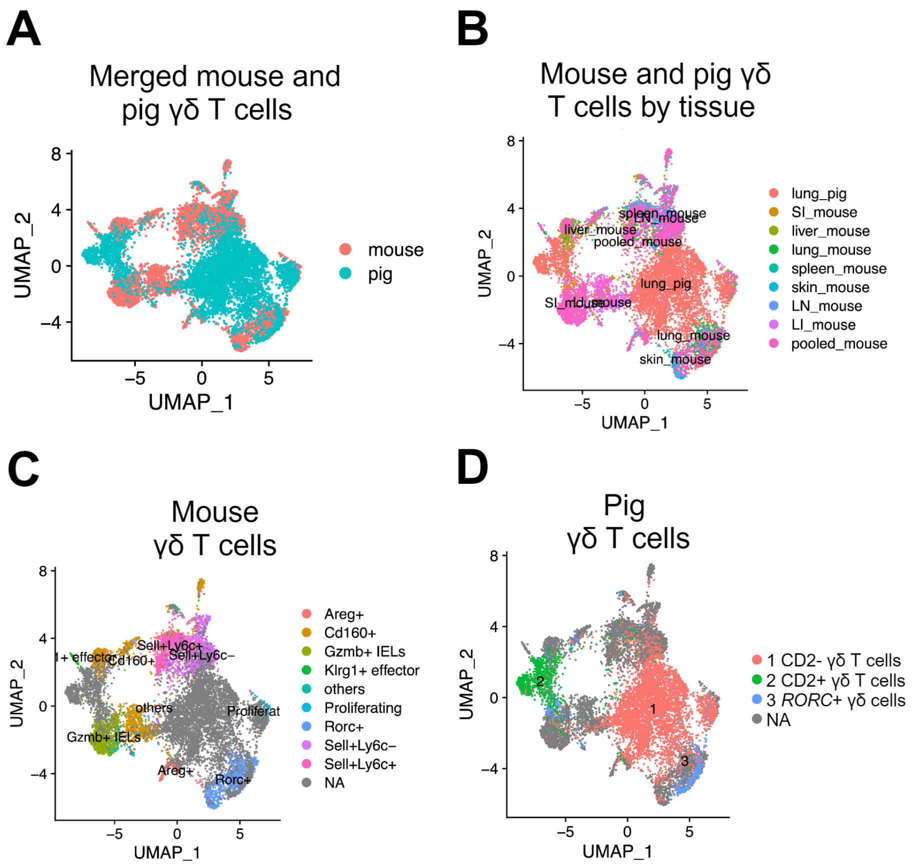

### Supplementary figure 3

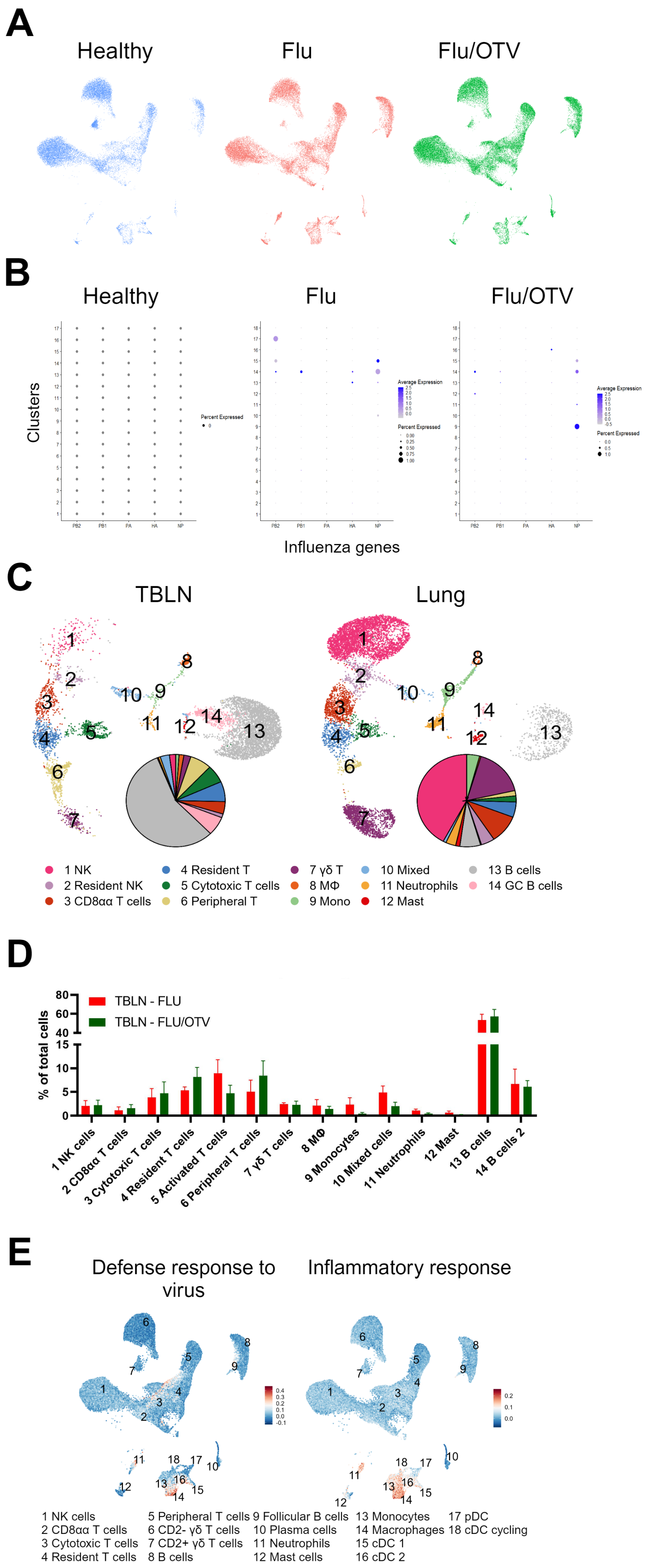

### Supplementary figure 4

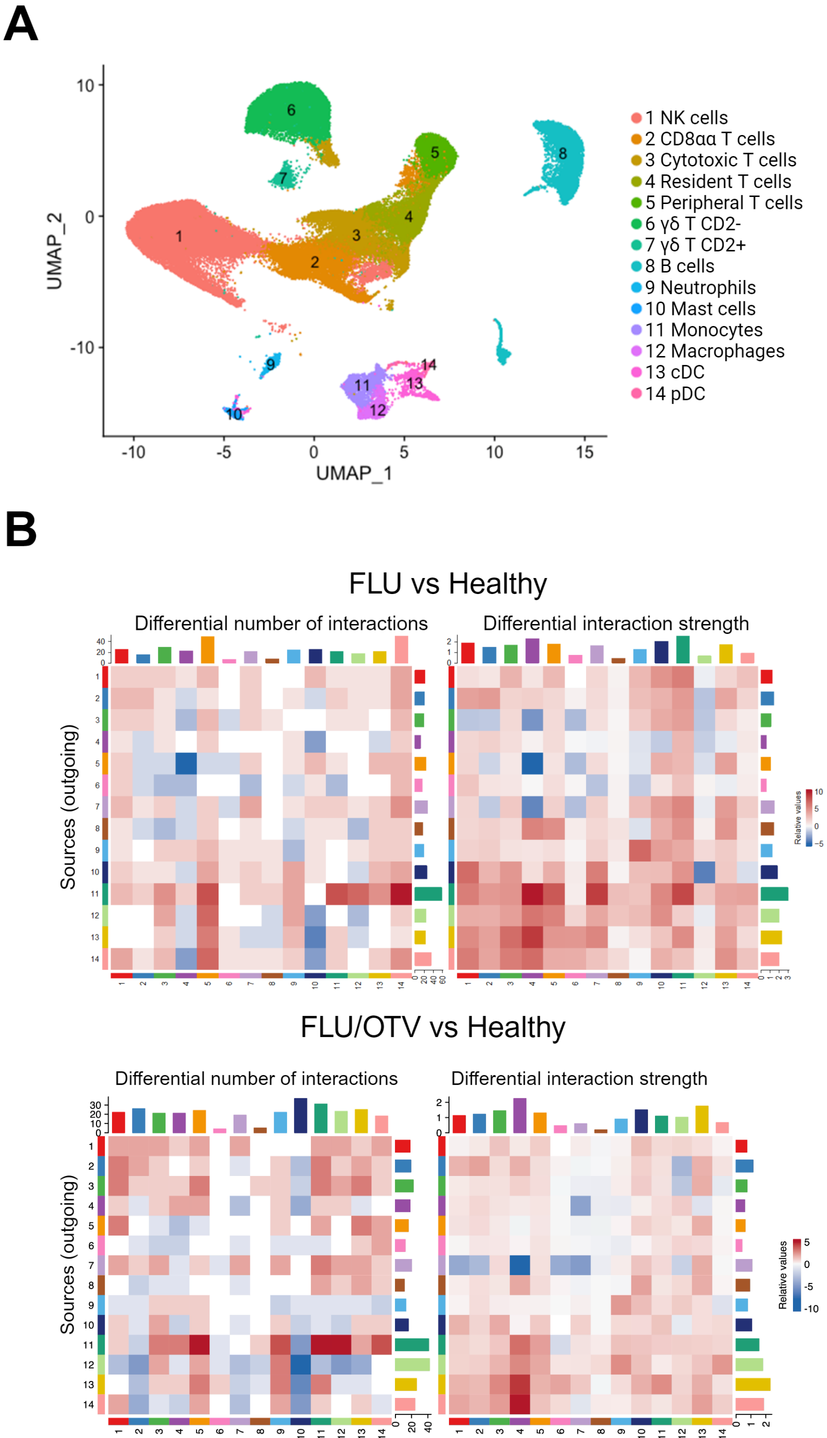

### Supplementary figure 5

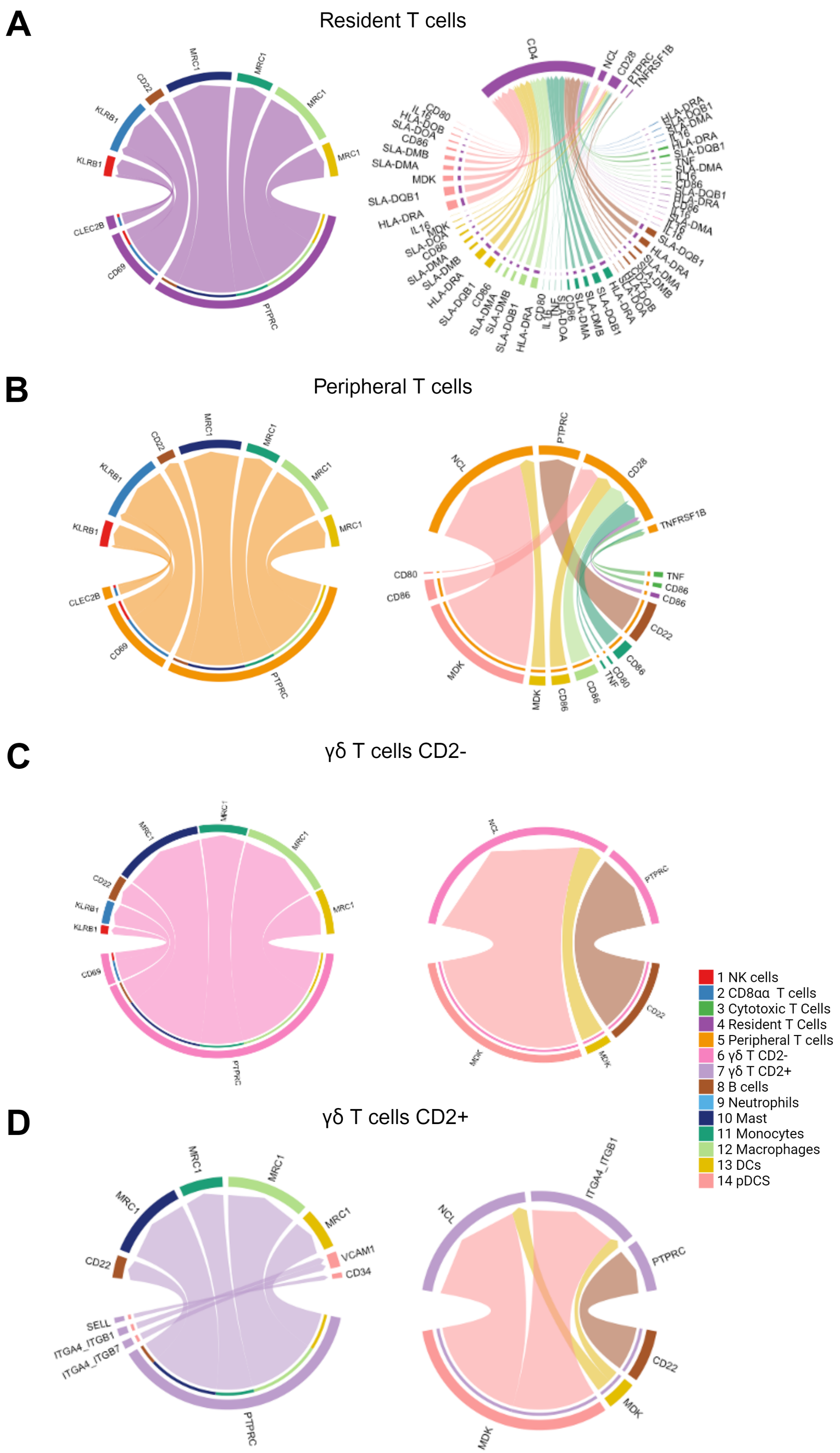

**E** Monocytes

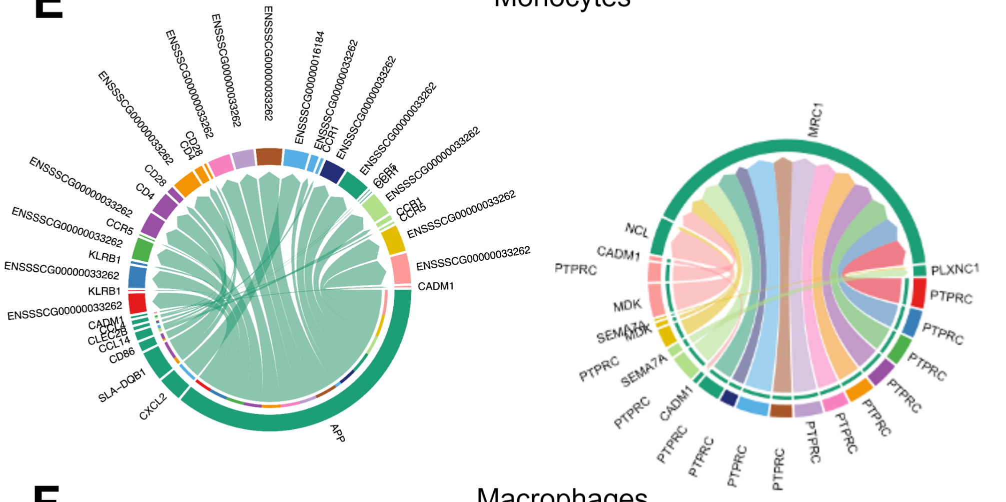


**F** Macrophages

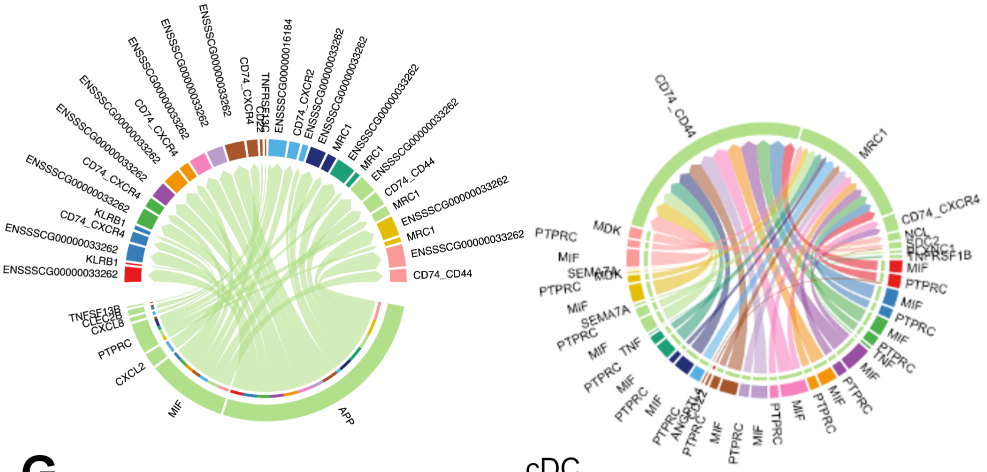


**G** cDC

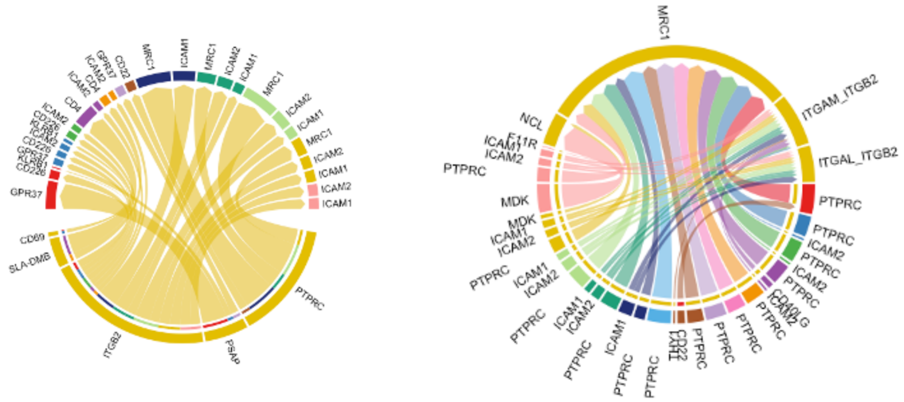


**H** pDC

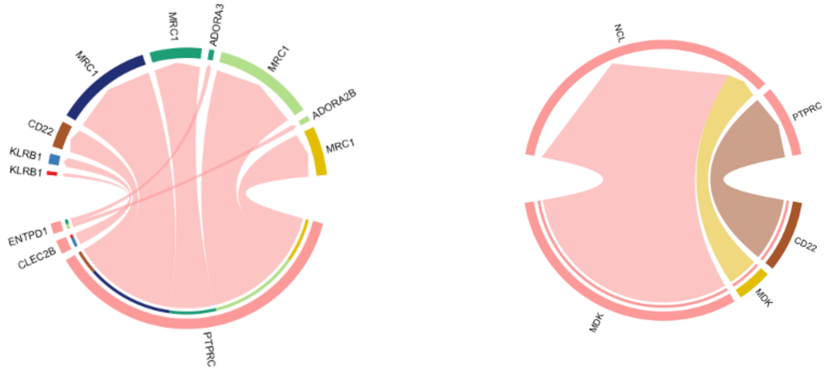

### Supplementary figure 6

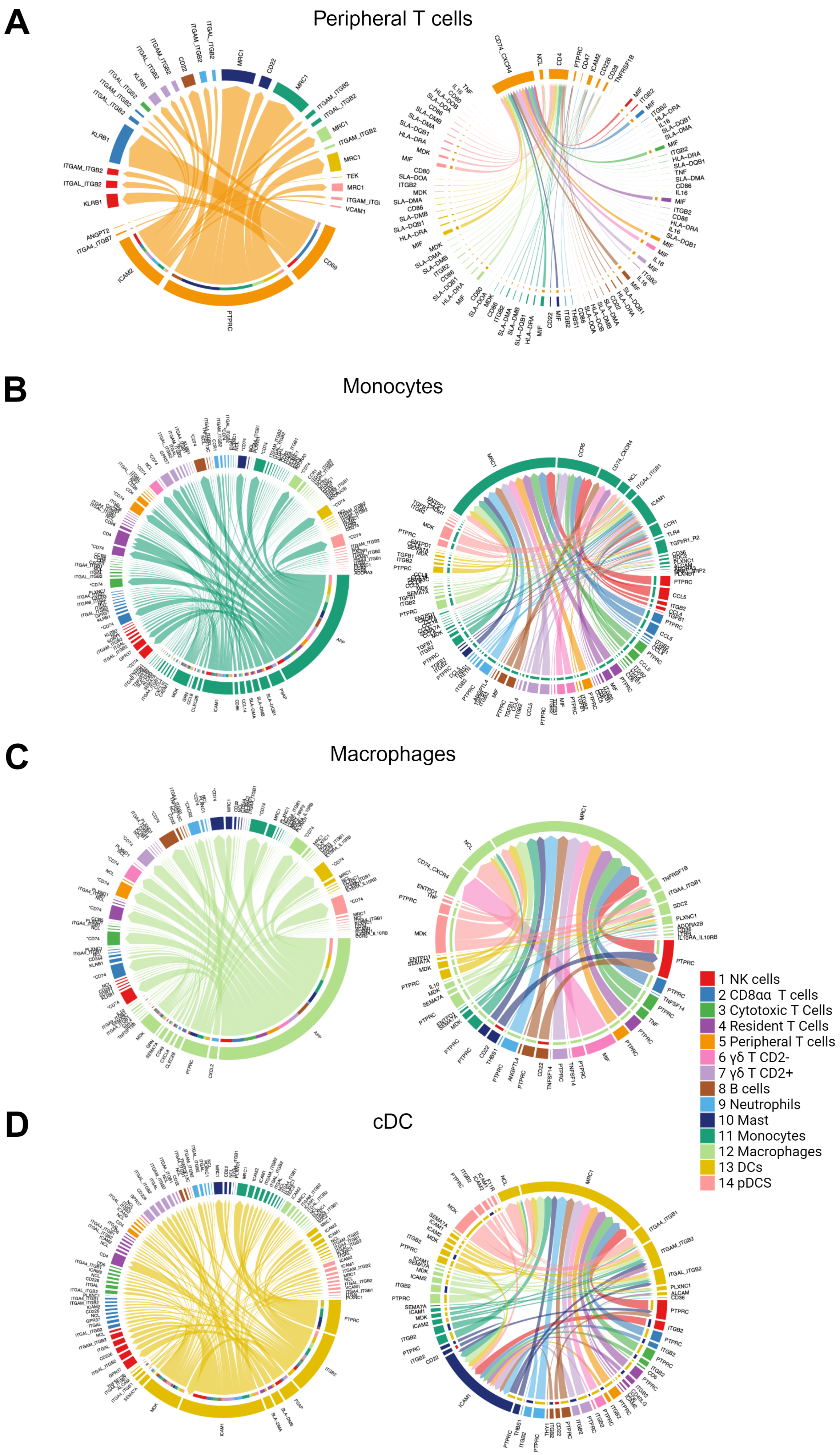
